# Substrate mediated mechanical forces enable optimal kinetic proofreading by T-cell receptors

**DOI:** 10.64898/2026.05.12.724610

**Authors:** Nicholas Jeffreys, Joshua M. Brockman, Tiam Heydari, Bryan A. Nerger, Wei-Hung Jung, Peter W. Zandstra, L. Mahadevan, David J. Mooney, Suraj Shankar

**Author notes:** Correspondence: L. Mahadevan, David J. Mooney, and S. Shankar. These authors contributed equally to this work (Nicholas Jeffreys, Suraj Shankar).

## Abstract

T-cells use molecular reactions with nonequilibrium error correction, i.e., proofreading, to discriminate between nearly identical antigens with high specificity and sensitivity. These receptor binding events are known to be force sensitive, yet traditional schemes of proofreading focus on reaction kinetics alone and do not consider the role of force dependent catch/slip bond behavior or interactions with mechanically engaged coreceptors such as adhesion molecules. To address this, we propose a minimal framework for proofreading of ligand discrimination by T-cell receptors (TCRs) that uses endogenous TCR mechanosensation and substrate-mediated mechanical interactions with adhesive proteins (load sharing) to improve recognition fidelity. We leverage the catch bond behavior of cognate antigens to delay decision making and amplify TCR signaling while discarding noncognate slip bond ligands in the presence of a force. By integrating our model with existing structural and molecular data, we show that substrate mechanics regulates the transmission of active cytoskeletal forces through a molecular clutch and controls the energization of bound TCRs needed for optimal proofreading. Our work demonstrates how mechanical forces and substrate properties can augment kinetic proofreading in T-cells, suggesting biomaterial design strategies for immunotherapies that tune the mechanical microenvironment of T-cells to achieve high fidelity TCR-ligand discrimination, antigen recognition, and activation.

## Main

The immune system recognizes and discriminates between self and foreign antigens. T-cells perform this task by binding T-cell receptors (TCRs) to peptide-loaded major histocompatibility complexes (pMHCs) decorated on antigen presenting cell (APC) surfaces. That TCRs can detect rare cognate pMHCs in a vast array of noncognate pMHCs with exceptionally high specificity and sensitivity is known to require a mechanism of nonequilibrium error correction^1–4^. Kinetic proofreading models have been suggested to explain this behavior by considering the activation of a TCR signaling cascade that expends energy to facilitate a sequence of kinetic delays^1-5^. During these delays, the progressive phosphorylation of available immunoreceptor tyrosine-based activation motifs (ITAMs) within the TCR/CD3-ζζ complex kinetically competes with the dissociation of the TCR-pMHC bond^5,6^. Consequently, transient interactions between TCRs and noncognate pMHC (n-pMHC) are unlikely to trigger T-cell activation, whereas longer-lived interactions between TCRs and cognate pMHC (c-pMHC) are more likely to support the formation of stable signaling complexes that enable T-cell activation (e.g., the LAT signalosome)^7,8^.

Complementing this biochemical picture, in recent years it has been suggested that physical forces can also support enhanced proofreading by dissipation of mechanical rather than chemical energy^9–13^. In-vitro experiments show that mechanically loaded TCR-pMHC complexes display force-dependent kinetics that can modulate proofreading^20,21^, through catch bond formation (with c-pMHC)^14,15^, slip bond formation (with n-pMHC)^16^, and T-cell adhesion to APCs (via LFA-1-ICAM-1 catch bonds)^17,18^. Molecular dynamics simulations of TCR-pMHC bond engagement and TCR mechanical allostery additionally support these experimental observations^9–11,19^. These studies largely focus on a fixed applied force, resulting in a constant mechanical energy budget for proofreading^13,15-17^. But cells can adaptively regulate cytoskeletal force generation through feedback to tune antigen discrimination, a feature that is often neglected; recent work in B-cells are an exception^20,21^. Furthermore, it remains unclear how active forces are transmitted to TCRs. Direct loading by motors is unlikely as TCRs are only weakly coupled to the actin cytoskeleton^22,23^, lacking direct structural connections^24^. This suggests that indirect force transmission through the environment is necessary for triggering mechanosensitive effects. While recent studies have highlighted how substrate properties can influence TCR discrimination efficacy^25,26^, an integrated framework for proofreading that couples cytoskeletal force generation, molecular kinetics, and substrate mechanics remains lacking.

We address this challenge by developing a minimal model for mechanically regulated proofreading that combines the reaction-limited kinetics of TCR-ligand discrimination with a molecular clutch description of LFA-1 adhesion to an elastic substrate exposing ICAM-1 ligands, rare c-pMHCs, or a large array of n-pMHCs. By incorporating experimentally known structural and molecular constraints, our framework captures key features of T-cell/APC contact mechanics during T-cell antigen discrimination, and incorporates catch/slip behavior exhibited by TCRs and LFA-1 coreceptors. Active forces are self-regulated by the molecular clutch and transmitted through the substrate, enabling mechanical cooperation between TCR-ligand and LFA-1 adhesion kinetics. Numerical simulations of the steady-state behavior show that resistance supplied by the substrate stiffness modulates the magnitude of piconewton (pN)-level loads generated by the LFA-1 clutch, and supplies TCR-pMHC bonds with mechanical energy during ligand discrimination. While extremely floppy or stiff APC surfaces abrogate TCR signal amplification by rare c-pMHCs, soft APC surfaces with intermediate stiffness convey physiological pN-loads to TCR-pMHC bonds, allowing optimal discrimination. Unlike conventional kinetic proofreading (where the amount of free energy that is expended to improve precision is fixed a priori in a speed/energy dissipation tradeoff), ‘mechanical’ proofreading depends on the mechanical properties of the T-cell microenvironment, such that energy transport to the TCR is dictated by mechanical filtering with LFA-1-generated forces against the APC substrate through actomyosin contractility.

Additionally, in the context of mechanical proofreading, the physical basis for discriminating between cognate and noncognate antigens hinges on the force-dependent kinetics of the TCR. Specifically, high-fidelity discrimination requires a maximal bond lifetime at a finite force that is characteristic of the catch/slip transition. The ratio of ‘correct’ to ‘incorrect’ signaling TCR-pMHC bonds (and by proxy, force-dependent bond lifetimes) yields a proofreading precision that distinctly peaks at this finite transition force. A fundamental question naturally arises: what dictates the magnitude of this applied force in the immunological synapse? Our model reveals that this optimal force is governed not solely by internal actomyosin contractility, but by the interplay between the substrate stiffness and the intrinsic rigidity of the T-cell membrane. In the extreme limits of substrate compliance—approaching either zero or infinite stiffness—the forces exerted on the TCR are generated almost exclusively by membrane mechanics rather than actomyosin motor activity through LFA-1. Consequently, a critical boundary condition for successful mechanical proofreading is that the force required to reach the catch/slip transition must be smaller than the maximal restoring force capable of being generated by the deformed membrane. Our results emphasize the potential mechanical determinants of TCR signal amplification, with implications for *in vitro* T-cell-material interface design and future mechanoimmunological investigation of TCR-ligand discrimination, antigen recognition, and activation.

Our mathematical framework for mechanical proofreading couples three basic modules:

i. TCR-ligand discrimination of c- or n-pMHCs through ITAM phosphorylation based proofreading steps.
ii. kinetics of an LFA-1-ICAM-1 molecular clutch^12,27^.
iii. T-cell adhesion-surface mechanics^28,29^.

### Reaction-limited, steady-state mechanokinetics of TCR-ligand discrimination with limited signaling **(i)**

We model TCR-ligand binding as a bimolecular reaction with distinct force dependent kinetic rates for cognate and noncognate ligands. Consistent with experimental measurements, we parametrize the TCR dissociation rate from c-pMHCs 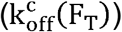 using a two-pathway model to capture a transition from catch to slip behavior (**eq. 1**)^30^, but instead use a simple Bell model for the dissociation rate from n-pMHCs 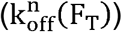, which only display slip bonding (**eq. 2**)^31^:

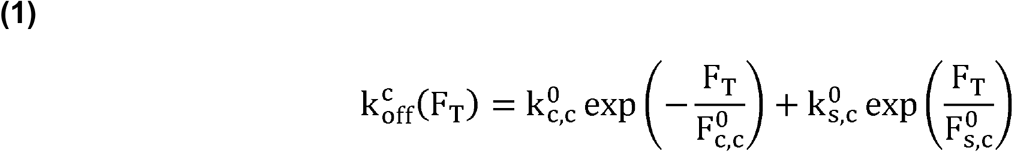

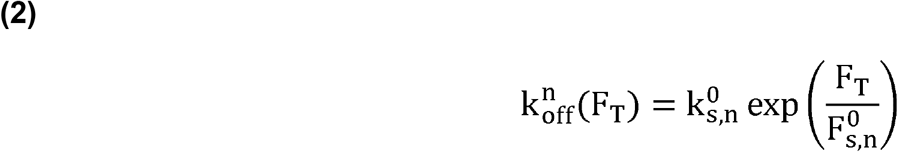

Here, 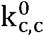 and 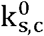are the catch/slip rates for TCR-c-pMHC unbinding at zero load, 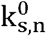is the slip rate for TCR-n-pMHC unbinding at zero load, and 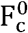 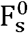are the force thresholds for the catch and slip bonds, respectively. We choose exponential forms for ease though there is emerging evidence that the bond lifetime distributions may be fat-tailed^32,33^. The force F_T_ experienced by a single bound TCR to either c or n-pMHC presented from the surface is modeled as a Hookean spring (**eq. 3**):

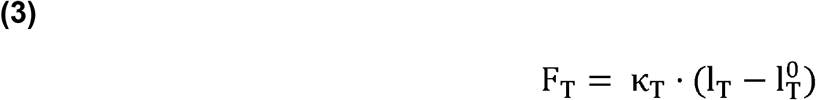

in which 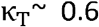 pN/nm is the TCR-pMHC bond stiffness^34^, l_T_ is the total length of the TCR-pMHC bond, and 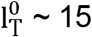nm is the TCR-pMHC rest length^35^.

Assuming mass conservation for the total number of receptors and ligands, we can solve the mass action kinetic equations (see Appendix for details) at steady-state to obtain the surface density of TCRs bound to either c or n-pMHCs (Π_c_ or Π_n_) to be (**eq. 4** and **eq. 5**):

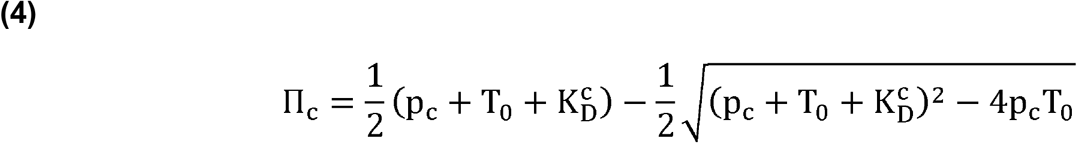

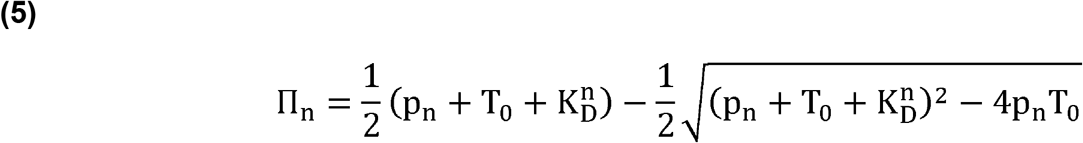

where T_0_ is the T-cell membrane density of available TCRs in the contact area (reacting-limiting), p_c_ (p_n_) are the dosed densities of c-pMHCs (n-pMHCs) presented on the surface, and 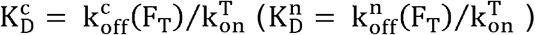 are the force-dependent dissociation constants of TCR-c-pMHC (TCR-n-pMHC) bonds.

Kinetic proofreading is enabled by irreversible phosphorylation of bound TCRs. After θ consecutive proofreading steps, the steady-state solution yields the density of activated, stable TCR-c-pMHC signaling complexes 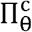 (**eq. 6**) or aberrant TCR-n-pMHC signaling complexes 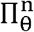 (**eq. 7**) to be (see Appendix for details),

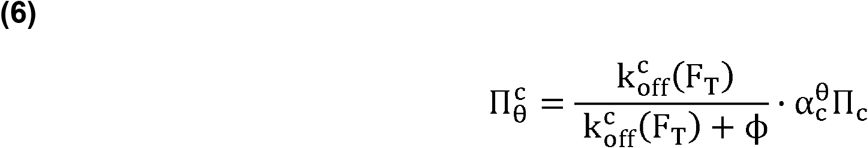

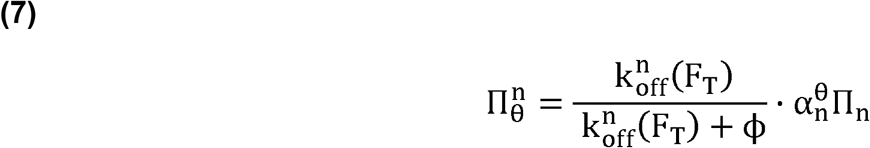

where ϕ is the limited signaling decay rate for rendering activated signaling complexes as non-signaling, and the ratios, 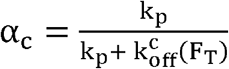 and 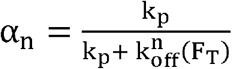capture the kinetic competition between the sequential tyrosine phosphorylation rate k_p_, and the pMHC unbinding rate during each proofreading step.

### Precision in TCR signal amplification and proofreading error rate

To quantify proofreading accuracy, we assume TCR signal amplification (denoted by R) is proportional to the density of activated signaling complexes, namely 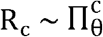 and 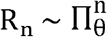(Eqs. **(6)** and **(7)**) for cognate and non-cognate ligands, respectively. The ratio of correct versus aberrant signal amplification then provides a measure of proofreading precision λ_a_ = Rc ⁄R_n_ (**eq. 10**):

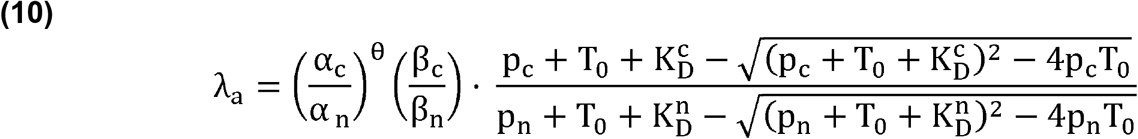

where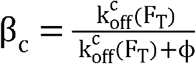 and 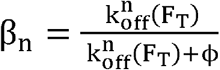 capture the kinetic competition between the limited signaling decay rate ϕ, and the pMHC unbinding rates. The inverse of λ_a_ gives a proofreading error rate f_0_ = 1/λ_a_.

### Molecular clutch dynamics of LFA-1-ICAM-1-mediated surface adhesion **(ii)**

Dynamics of active surface adhesion is modeled as a molecular clutch, wherein the reversible binding of LFA-1 coreceptors to ICAM-1 is described using first order kinetics (**eq. 14**):

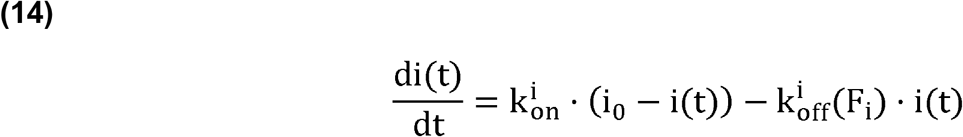

Here, i(t) is the surface density of bound LFA-1-ICAM-1 complexes, i_0_ is the surface density of LFA-1 coreceptors, 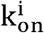 is the binding rate, and 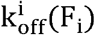 is the force-dependent unbinding rate. The force exerted by a single bound adhesion complex is 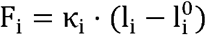, where l_i_ is the total bond length, κ_i_ ∼ 0.2 pN/nm the bond stiffness^36^, and 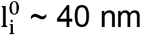 the rest length^35^. Catch bond behavior of integrins is incorporated in the unbinding rate 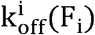, parametrized by a two-pathway model (**eq. 15**):

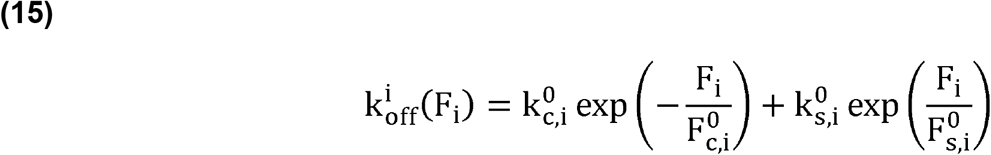

that transitions to slip bonding at large loads. The zero load catch and slip rates are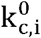 and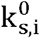, along with the corresponding transition forces, 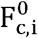 and 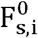. When bound, the LFA-1-ICAM-1 adhesion complex engages with the actomyosin cytoskeleton and is stretched by motor activity, but this elastic extension y_i_ is relaxed upon unbinding^37^. Using a mean-field approach that averages over stochastic binding events (see Appendix for derivation), we obtain the dynamics of the average stretch ⟨y_i_⟩ to be (**eq. 16**):

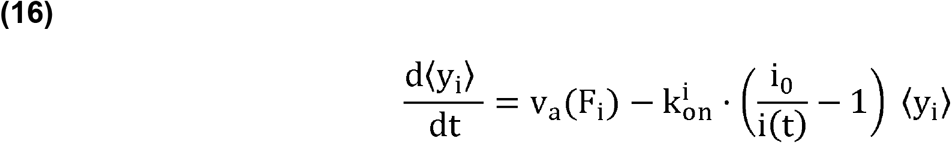

where v_a_(F_i_), the retrograde flow velocity of the actomyosin gel obeys a simple linear force-retrograde flow velocity relationship (**eq. 17**):

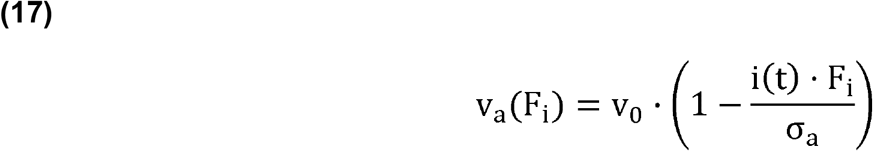

with v_0_ ∼ 60 nm/s being the maximal (zero load) retrograde flow velocity, and σ_a_ ∼ 1 kPa being the contractile actomyosin stall stress^38,39^, beyond which actin flows are arrested.

At steady-state, the kinetics and kinematics of LFA-1-mediated adhesion can be solved to obtain (**eq. 18** and **eq. 19**):

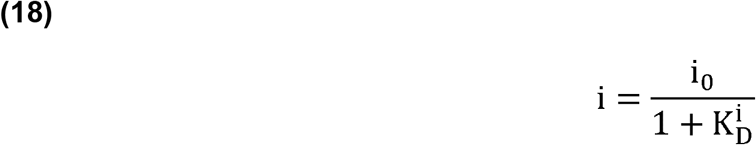

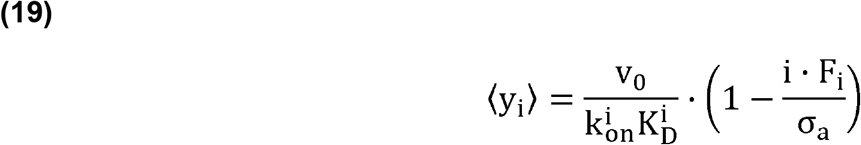

where 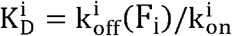is the force dependent equilibrium dissociation constant. Eq. 19 encodes mechanical feedback in active force generation and can be simply understood by noting that the) to the total length of the adhesion complex stretches at speed v_a_ for a bond lifetime 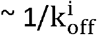 yielding an average extension 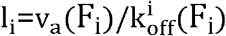. To relate this actively generated intracellular stretch (⟨y_i_⟩) to the total length of the adhesion complex (l_i_), we need to specify the kinematics of the substrate, as done below.

### T-cell adhesion-surface mechanics **(iii)**

We complete the model by imposing force balance to mechanically couple TCR-antigen kinetics (i) with the active dynamics of adhesion (ii). In the absence of direct links to the cytoskeleton, bound TCRs can stretch and generate forces only when resisted by the T-cell membrane. Previous models of synaptic patterning^23,40,41^ have shown how differential molecular size (or receptor and adhesive proteins) coupled to membrane elasticity can generate spatially segregated TCR clusters. Here, we neglect this spatial patterning dynamics and instead focus on an effective description that presumes the existence of preformed clusters. Assuming the T-cell and opposing APC are separated by an average gap height h ∼ 50-60 nm due to large glycosylated proteins (e.g., CD45 ectodomain^42^), we consider TCR-antigen bonds to deform the membrane locally (displacement δh) on the scale of a TCR microcluster^43–45^, R ∼ 50 nm^46^. On the other hand, elastic deformations of the APC surface (u) balance the shared load exerted by both receptor and adhesion molecules, extended across the contact area, A_c_ ∼ 3 × 10^4^ nm^2^, of say a microvillus^47^. Altogether, force balance at the T-cell membrane and the APC surface then yields (**eq. 20** and **eq. 21**):

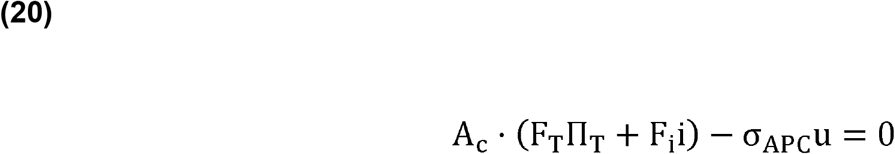

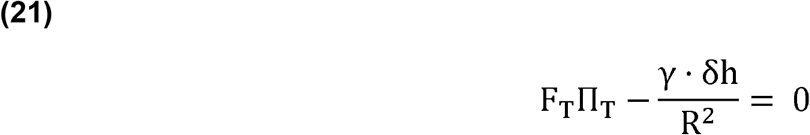

where σ_APC_ is the APC surface stiffness and γ ∼ 0.03 pN/nm is the T-cell membrane tension. The total bond lengths that enable force generation on TCR-pMHC (l_T_) and LFA-1-ICAM-1 (l_i_) complexes are related to the displacements of the APC surface (u), T-cell membrane (δh), and the intracellular bond stretch of LFA-1-ICAM-1 ⟨y_i_ ⟩ as follows:

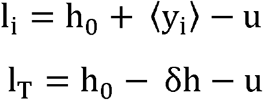

These relations encode structural and geometric constraints intrinsic to the T-cell adhesive contact and complete our model formulation.

## Results

### The magnitude of receptor-ligand force modulates reaction rates in the mechanical proofreading framework

As a first step, we ask: how do receptor-ligand forces impact kinetic proofreading? Kinetic proofreading models with limited signaling have been shown to be consistent with the majority of published experimental data^1^. We incorporate force dependence within this framework simply via the force-sensitive unbinding rates (**eq. 1-2, eq. 15**), see **Fig. 1a-1d**. For simplicity, here we directly specify the force on each bond and neglect substrate mediated mechanical couplings. In **Fig. 1e**, we plot the force-sensitive bond lifetimes (1⁄k_off_(F)) and dissociation constants (K_D_) against physiological pN-level loads for human OT1(†) TCRs binding either c-pMHCs (OVA antigen on H2-Kb MHC class I) or n-pMHCs (R4 antigen on H2-Kb MHC class I), and LFA-1 binding ICAM-1 (in the presence of Ca^2+^, Mg^2+^, and CXCL12) using parameters extracted from 2-dimensional micropipette aspiration experiments (**Appendix Table a1**)^30,31^. The maximal bond lifetimes for TCR-c-pMHC and LFA-1-ICAM-1 center around 8-12pN of load, while TCR-n-pMHC exhibits an exponential decaying bond lifetime under increasing load^14,17^. To quantify relative competition in unbinding, we examine the ratios of off rates as a function of a common force (**Fig. 1f**). Upon increasing the load, TCR-n-pMHC off rates dominate over that of TCR-c-pMHC, whereas LFA-1-ICAM-1 off rates always exceeded the rates of TCR-antigen unbinding (cognate and noncognate). This is simply understood by noting that cognate antigens form catch bonds with longer lifetimes at high load, in contrast to short lived slip bonds formed by noncognate antigens.

**Fig. 1.**
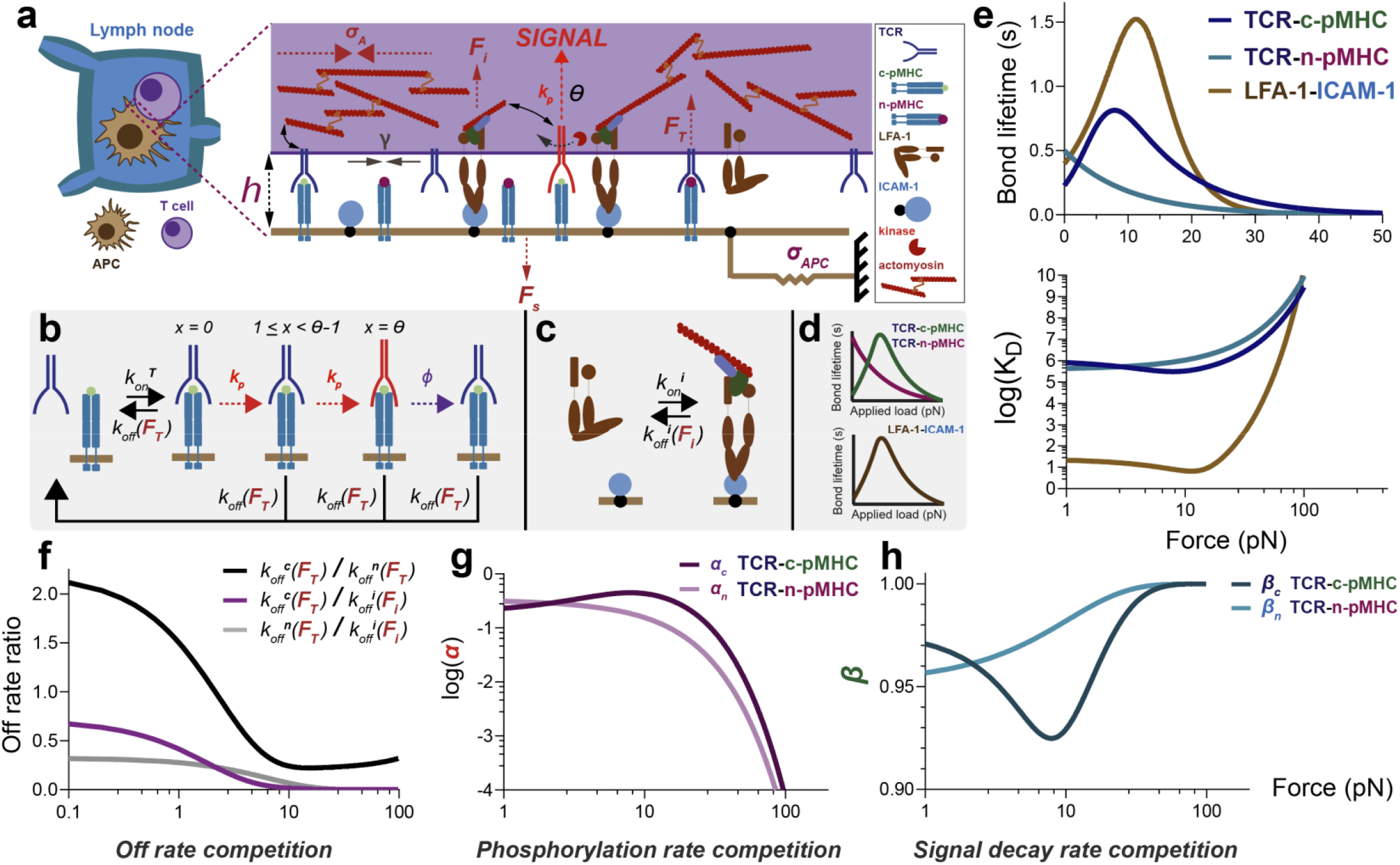
The magnitude of receptor-ligand force dictates reaction rate competition in the mechanical proofreading framework. a) Schematic of mechanical proofreading model recapitulating key features of a T-cell adhering to an elastic surface decorated with c-pMHC, n-pMHC, and ICAM-1. b) Mechanical proofreading framework that incorporates limited signaling and catch/slip bond behavior exhibited by TCR-ligand binding. c) Kinetics and kinematics of LFA-1-ICAM-1 bond formation in a molecular clutch framework. Force generation in response to resistance supplied by the elastic surface facilitates LFA-1-mediated adhesion, and passively energizes TCR-ligand binding through the injection of mechanical energy by force balance. d) TCR-c-pMHC and LFA-1-ICAM-1 bonds exhibit catch bond behavior, whereas TCR-n-pMHC bonds exhibit slip bond behavior. e) Bond lifetimes and dissociation constants of TCR-c-pMHC, TCR-n-pMHC, and LFA-1-ICAM-1 binding kinetics as a function of force. f) Off rate competition between TCR catch/slip and LFA-1 catch bond off rates. g) ITAM phosphorylation rate competition with TCR-c-pMHC catch bonds and TCR-n-pMHC slip bonds. h) Limited signaling complex decay rate competition with TCR-c-pMHC catch bonds and TCR-n-pMHC slip bonds.

Mechanical forces also influence the proportion of bound TCRs that survive TCR-ITAM phosphorylation by tyrosine kinases (e.g., ZAP-70^48^) during a proofreading step. We find that the phosphorylation rate for TCR-c-pMHCs (quantified by α_c_) dominates over the off rate around 8-10pN, whereas relative phosphatase activity decreases at loads greater than 15pN (Fig. 1g). In contrast, the phosphatase activity for TCR-n-pMHCs (α_n_) continually decreases with respect to increasing load (**Fig. 1g**). In addition, both TCR-c-pMHCs and TCR-n-pMHCs can form stable signaling complexes post-proofreading that can be rendered non-signaling and dissociated due to a signal decay rate ϕ (**Fig. 1b**). We quantify this kinetic competition using β_c_ and β_n_ and plot their dependence on the force in **Fig. 1h**. For signaling TCR-c-pMHCs under 8-10pN of load, the signal decay rate dominates over the off rate, whereas signaling TCR-n-pMHCs have a larger unbinding rate compared to signal decay.

### The magnitude of receptor-ligand force controls TCR-ligand proofreading precision and error

To assess how proofreading is impacted by forces, we next compute the steady-state densities for the cognate and noncognate ligands. (**eq. 6-10**). We choose the n-pMHC surface density (p_n_ = 2·10^4^ μm^-2^) to be 10-fold higher than the c-pMHC and TCR densities (p_C_,T_0_= 2·10^3^ μm^-2^) to mimic discrimination of rare cognate antigen from noncognate antigen decorating APC surfaces (**Appendix Table a1**)^47^. Despite this difference in density, cognate antigens form long-lived catch bonds in contrast to short-lived slip bonds formed by noncognate antigens. So we expect an applied force can still enhance TCR-ligand discrimination by minimizing the formation of erroneous TCR-n-pMHC signaling complexes, while maximizing the formation of ‘correct’ TCR-c-pMHC signaling complexes (**Fig. 2a**). **Fig. 2b** shows the impact of receptor-ligand force on the formation of signaling TCR-c-pMHC and TCR-n-pMHC complexes (**eq. 6-7**) across θ proofreading sequences. Here, θ is set to 10 as the TCR uniquely contains 10 ITAMs to transduce pMHC binding into downstream signaling cascades (**Appendix Table a1**)^49^. Corresponding to the optimal TCR-c-pMHC bond lifetime, we find that the maximal formation of stable, signaling TCR-c-pMHC complexes is centered around an applied load F ∼ 8-10pN (**Fig. 2b**). Conversely, force application and proofreading both suppress the formation of erroneous TCR-n-pMHC signaling complexes through rapid dissociation (**Fig. 2c**).

**Fig. 2.**
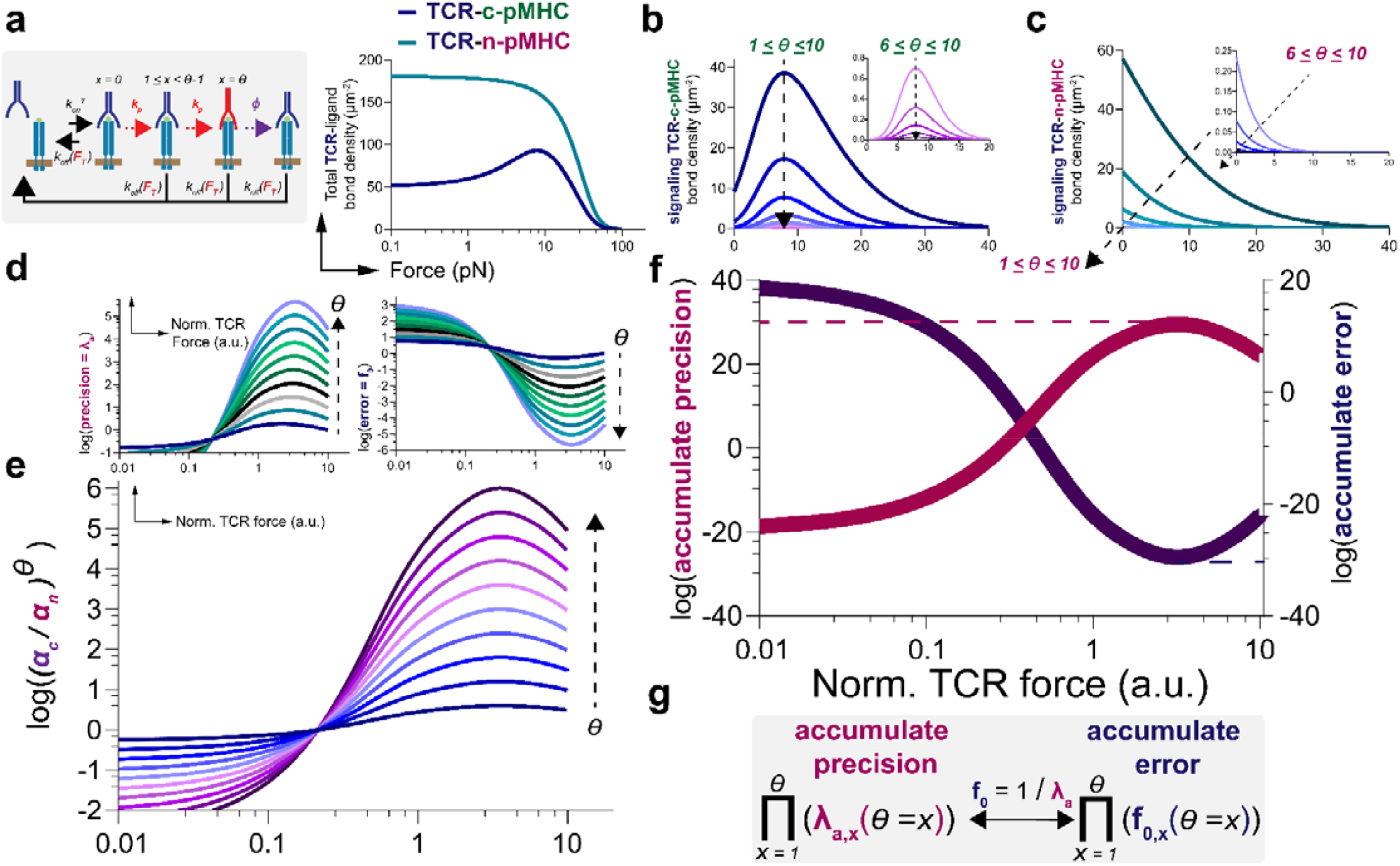
The magnitude of receptor-ligand force controls TCR-ligand proofreading precision and error. a) Schematic of mechanokinetic TCR-ligand discrimination with catch/slip off rates and TCR-ligand bond interfacial surface density as a function of force. b) Signaling competent TCR-c-pMHC bond interfacial surface density as a function of force from 1 < 0 < 10 proofreading sequences. c) Aberrant signaling competent TCR-n-pMHC bond interfacial surface density as a function of force from from 1 < 0 < 10 proofreading sequences. d) TCR signal amplification precision and error as a function of force from 1 < 0 < 10 proofreading sequences. e) Kinetic competition of c-pMHC and erroneous n-pMHC-mediated TCR phosphorylation as a function of mechanical force from 1 < 0 < 10 proofreading sequences. f) accumulate precision and error of TCR signal amplification as a function of mechanical force. g) accumulated error and precision are calculated by taking the total product of precision and error across 1 < 0 < 10 proofreading sequences. TCR force is normalized to 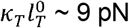.

Combining these results, we plot the precision (**eq. 10**) and error rate (**eq. 13**) of TCR signal amplification during proofreading as a function of the applied force (**Fig. 2d-f**). As expected, precision is increased with each proofreading sequence with optimal performance achieved at a load F ∼ 10 pN corresponding to the characteristic force scale at which cognate bonds transition from catch to slip behavior. This response is predominantly controlled by the proofreading metrics (α and α_n_, **eq. 8-9**) that quantify the kinetic competition between the sequential tyrosine phosphorylation rate k_p_, and the c or n-pMHC off rate during each proofreading step, as seen in **Fig. 2e**. Similar trends are also observed in the accumulated precision (or error) across all proofreading steps (**Fig. 2f, 2g**), consistent with recent results^13^.

### LFA-1 molecular clutch dynamics modulate the mechanics of LFA-1-ICAM-1 adhesion and TCR-ligand bond formation

We now relax the assumption of a constant force and investigate how actively regulated forces and substrate mediated mechanical cooperation between LFA-1 and TCR modulate their kinetics. As the TCR is not directly coupled to actomyosin, TCR force generation in the model is mediated by the LFA-1 molecular clutch through deformations of the T-cell membrane and adherent APC surface (**eq. 20-21**)^9,18,50,51^. TCRs binding to n-pMHC cause larger deformation and stress in the T-cell membrane, compared to c-pMHCs across varying APC surface stiffness (**Appendix Fig. 1a**), despite negligible difference in APC surface displacement between cognate and noncognate antigens (**Appendix Fig. 1b**). This suggests that membrane flexibility primarily controls the differential forces experienced by TCR-c-pMHC and TCR-n-pMHC bonds. Increasing APC stiffness > 1 pN/nm causes the actomyosin retrograde flow to stall (**Appendix Fig. 2a**) as the density of LFA-1-ICAM-1 bonds saturates (**Appendix Fig. 2b**). Correspondingly, the forces generated by LFA-1, TCRs engaged with c-pMHC or n-pMHC, all follow a sigmoidal increase with APC stiffness, with values within their physiologically measured ranges (**Appendix Fig. 2a**)^14,18^. In other words, the APC surface mechanically gates the transmitted force, switching on the maximum load on receptor antigen bonds beyond a characteristic stiffness ∼ 1 pN/nm, when the motor (actomyosin flow) stalls.

The bond lifetimes and dissociation constants for TCR-c-pMHCs, TCR-n-pMHCs, and LFA-1-ICAM-1 complexes vary with respect to the APC stiffness through the force they respectively experience (**Appendix Fig. 2c**). LFA-1-ICAM-1 bond lifetimes increases with APC surface stiffness, plateauing at ∼1.5 seconds, consistent with the stalling of actomyosin observed and the formation of LFA-1-ICAM-1 adhesion complexes (**Appendix Fig. 2a, 2b**). TCR-c-pMHC bond lifetime instead has a biphasic dependency on APC surface stiffness, reflecting the transition from catch to slip behavior, with the maximal lifetime achieved for a stiffness ∼ 1 pN/nm. Conversely, TCR-n-pMHC bond lifetimes decay as surface stiffness is increased. As a result, the formation of TCR-c-pMHC bonds is increased around an APC surface stiffness of 1 pN/nm, but TCR-n-pMHC slip bonds simply dissociate as APC surface stiffness approaches supraphysiological membrane rigidities (e.g., 1000 pN/nm) (**Appendix Fig. 2d**).

How does this response depend on the extent of catch or slip bonding behavior? As an example, we consider human OT1(†) TCRs binding c-pMHCs of varying catch bond amplitude (OVA, A2, G4, or E1 loaded on H2-Kb MHC class I, **Appendix Table a2**) and compare them against the TCR-n-pMHC (OT1(†) TCR binding R4 antigen loaded on H2-Kb MHC class I) slip bond as a function of receptor-ligand force and APC surface stiffness^14,52^. We compute the change in the Gibbs free energy of binding (ΔG^°^, **Appendix eq. a19**) to quantify the thermodynamics of TCR engagement. Consistent with our previous analysis of the TCR-pMHC dissociation constant (K_D_, **Fig. 1e** and **Appendix Fig. 2c**), we find that the largest decrease in free energy (in units of k_B_T) is achieved for OT1(†) TCR-OVA-MHC-I catch bonds at intermediate loads (force ∼ 8-12 pN, or APC stiffness ∼ 1 pN/nm; see **Fig. 3a-d**). In contrast, cognate antigens forming weaker catch bonds (e.g., E1, G4) minimize their change in Gibbs free in energy at zero load or low APC stiffness (**Fig. 3a, 3c**) and do not benefit from mechanical forces in improving TCR engagement. This implies that the amplitude/magnitude of the TCR-ligand catch bond, not just its presence, crucially matters in determining whether mechanical forces can induce a spontaneous thermodynamic process^53^ allowing TCR signal amplification against a slip-bonding n-pMHC.

**Fig. 3.**
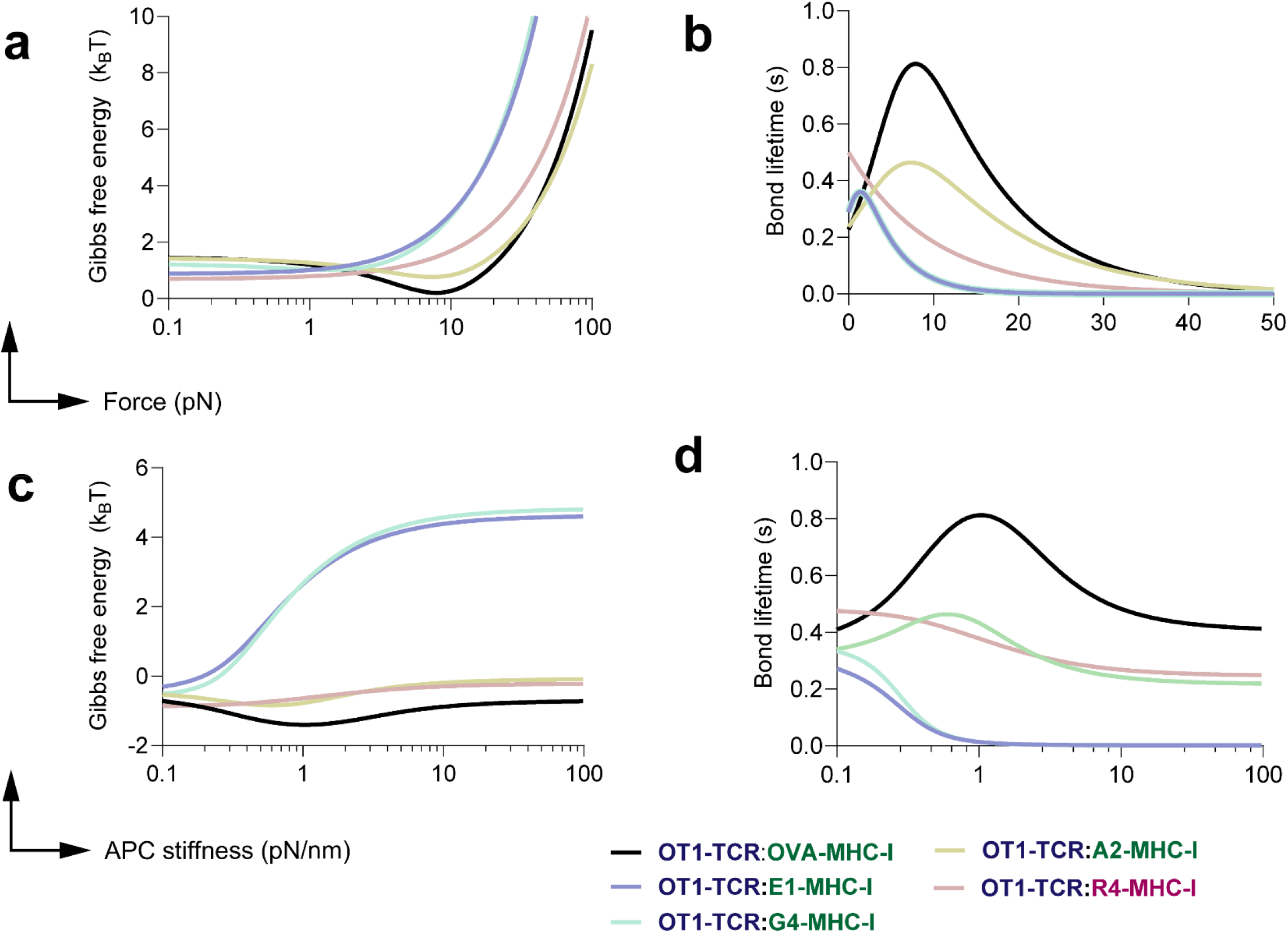
TCR-pMHC catch/slip bond amplitude and Gibbs free energy minimization are controlled by receptor-ligand force magnitude and APC surface stiffness. a) Standard state change in Gibbs free energy of OT1(†) TCRs engaging various c-pMHCs or R4 n-pMHC as a function of force. b) Catch/slip bond behavior exhibited by OT1(†) TCR-ligand binding as a function of force. c) Standard state change in Gibbs free energy of OT1(†) TCRs engaging various c-pMHCs or R4 n-pMHC as a function of APC surface stiffness. d) Catch/slip bond behavior exhibited by OT1(†) TCR-ligand binding as a function of APC surface stiffness.

### Soft APC surfaces maximize TCR-ligand proofreading precision

We combine the above results to evaluate the impact of surface mechanics on TCR-ligand proofreading precision (**Fig. 4a**). **Figs. 4b, 4c** show the density of signaling TCR-c-pMHC and TCR-n-pMHC complexes across θ = 10 proofreading sequences as a function of APC surface stiffness. We find that soft APC surfaces with intermediate stiffness ∼ 1 pN/nm allow for the optimal TCR-c-pMHC bond lifetime accompanied by the maximal formation of stable, signaling TCR-c-pMHC complexes, whereas stiffnesses approaching 0 pN/nm or 1000 pN/nm leads to suboptimal signaling (**Fig. 4b**). This is additionally reflected in the suppressed formation of erroneous TCR-n-pMHC signaling complexes for stiff substrates (σ_APC_ ≥ 1 pN/nm), see **Fig. 4c**. As a result, we find that TCR proofreading precision is optimal for soft APC surfaces across all θ proofreading step (**Fig. 4d-e**), with a concomitant enhancement of the accumulated precision as well (**Fig. 4e**). Very low or high APC stiffness (> 1 pN/nm) leads to lower proofreading precision (and thus greater error), suggesting that the substrate serves as a mechanical filter that enables optimal performance when force generation by the molecular clutch is matched with force transmission through the environment. Finally, to test the dosage dependence of our results, we increase the c-pMHC dose to supraphysiological values and find the receptor-ligand force is diminished (due to load sharing), causing the optimal stiffness to shift to higher values (**Appendix Fig. 3a, 3b)**, consistent with experimental observations^9,54^. Eventually, for very high c-pMHC doses, the biphasic dependence of precision on stiffness is altogether abolished (**Appendix Fig. 3a, 3b)**. Similarly, decreasing n-pMHC dose enhances proofreading precision, transitioning from a monotonic to a biphasic response as well (**Appendix Fig. 3c, 3d**).

**Fig. 4.**
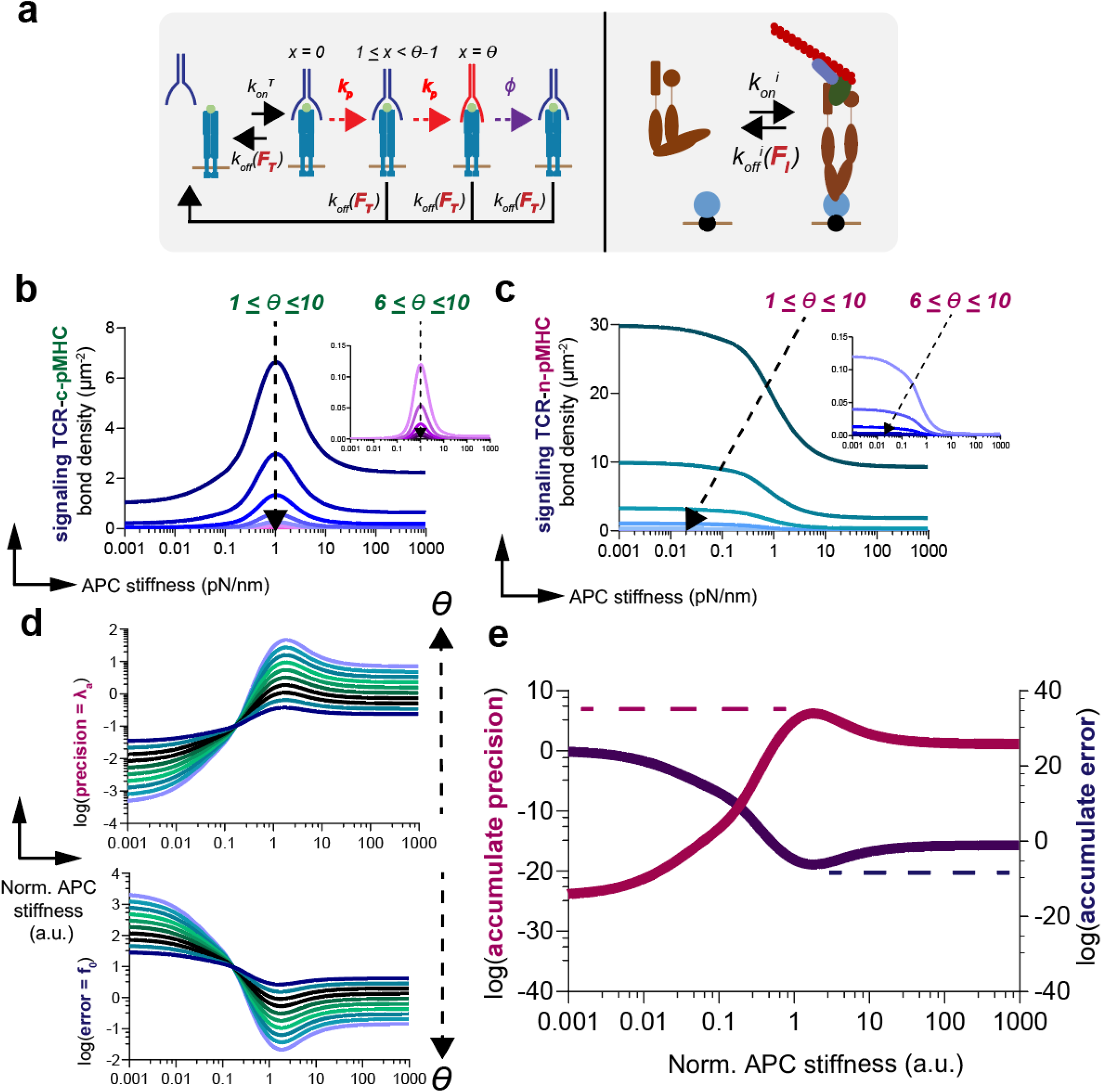
LFA-1 molecular clutch dynamics at steady-state modulates TCR-ligand proofreading precision and error in response to APC surface stiffness. a) Schematic of mechanokinetic TCR-ligand discrimination with catch/slip off rates in concert with LFA-1 molecular clutch dynamics. b) Signaling competent TCR-c-pMHC bond interfacial surface density as a function of APC surface stiffness from proofreading sequences. c) Aberrant signaling competent TCR-n-pMHC bond interfacial surface density as a function of APC surface stiffness from from proofreading sequences. d) TCR signal amplification precision and error as a function of APC surface stiffness from proofreading sequences. e) accumulate precision and error of TCR signal amplification as a function of APC surface stiffness. APC surface stiffness is normalized to 1 pN/nm.

### TCR signal amplification is modulated by energy transport from the active power generated by LFA-1-ICAM-1 adhesions

What are the energetic costs of this mechanism of proofreading? Conventional kinetic proofreading biases reactions to amplify discrimination accuracy by dissipating a fixed budget of chemical free energy^55–57^, as captured by the irreversible ITAM phosphorylation steps in our model. But here, reactions can also be biased by forces that are actively generated and regulated, leading to mechanical energy costs that we focus on. To assess how mechanical energy is partitioned during antigen discrimination, we separately evaluate the elastic energy stored in the APC surface 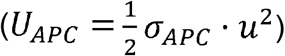, LFA-1-ICAM-1 adhesions 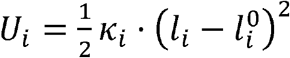, and TCR-pMHC bonds 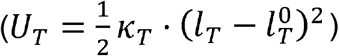. The nonequilibrium power (P) generated by the actomyosin cortex and dissipated through the adhesive dynamics of the molecular clutch is given by:

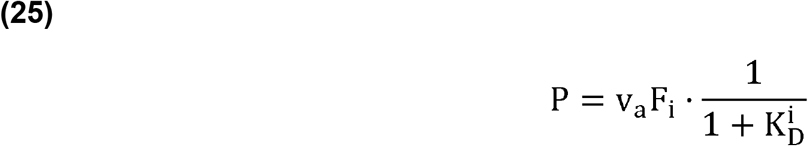

We first examine the elastic potential energy stored in human OT1(†) TCRs binding c-pMHCs of varying catch bond amplitude (OVA, A2, G4, or E1 loaded on H2-Kb MHC class I) and compare them against the TCR-n-pMHC (OT1(†) TCR binding R4 antigen loaded on H2-Kb MHC class I) slip bond as a function APC surface stiffness (in units of k_B_T, **Fig. 5a, eq. 22-24**). In all cases, we observe that increasing APC surface stiffness allows a larger amount of elastic energy to be stored in bound TCRs, with the relatively weak catch bonds, OT1(†) TCR-E1-MHC-I and OT1(†) TCR-G4-MHC-I, storing the most elastic energy. The active power dissipated by LFA-1 also increases with APC stiffness, and saturates beyond σ_APC_ ≥ 1 pN/nm, for the different TCRs considered. But in contrast to a traditional kinetic proofreading mechanism, we observe that greater nonequilibrium dissipation and energization of bonds actually leads to lower proofreading precision (**Fig. 5b, 5c, eq. 25**). Strikingly, OT1(†) TCRs binding the strong catch bond forming antigen OVA-MHC-I display enhanced proofreading precision with a comparable power expenditure for the cognate and non-cognate antigen (relative power = P_c_/P_n_ ∼ 1, **Fig. 5c**). In contrast, TCRs engaging with weak catch bond forming antigens (A2, G4, or E1) expend more than 3-fold the nonequilibrium power dissipated in the non-cognate case (OT1(†) TCRs binding R4 loaded MHC-I), only to have poorer discrimination (precision λ_a_ < 1), realizing a mechanical variant of an “anti-proofreading’’ regime^58^.

**Fig. 5.**
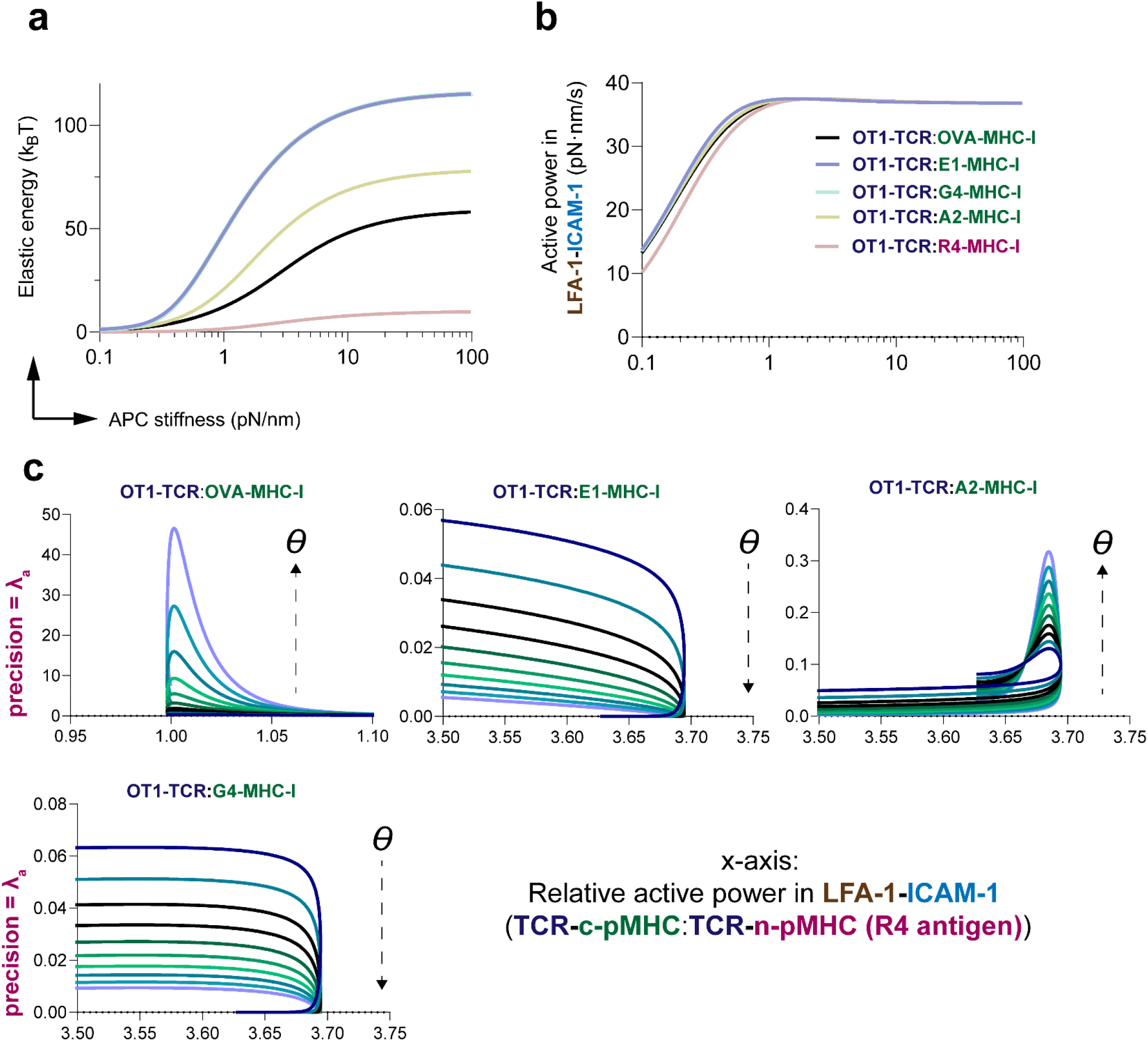
Active power generated by LFA-1-ICAM-1 bonds and TCR catch/slip bond amplitude enables TCR signal amplification through elastic energy storage in TCR-pMHC bonds. a) Elastic energy storage (in units of k_B_T) in TCR-pMHC bonds with different antigen types as a function of APC surface stiffness. b) Active power (energy transport) in total bound LFA-1-ICAM-1 complexes as a function of APC surface stiffness for OT1(†) TCRs engaging varying c-pMHC or n-pMHC. c) TCR signal amplification precision as a function of the relative ratio of active power in LFA-1-ICAM-1 complexes when OT1(†) TCRs engage different c-pMHC relative to R4-loaded n-pMHC.

### Mechanical filtering with LFA-1-ICAM-1 adhesions determines TCR signal amplification

How can we understand these results? Unlike conventional proofreading where energy is directly provided to the discriminating reaction, here, TCRs can harness nonequilibrium mechanical energy only indirectly, by transmission through substrate adhesion. This suggests the force generating motor (actomyosin and LFA-1-ICAM-1), the transmitter (substrate), and force receiver (TCR-antigen bond) must all be ‘impedance matched’, to use an electrical analogy, for optimal transfer of usable power for signal amplification. Without the optimal coupling, the nonequilibrium dissipation is futile and unavailable to aid proofreading. To quantify such impedance matching, we directly compare the stored elastic energy in the APC surface, along with the OT1(†) TCR-OVA-MHC-I (cognate), OT1(†) TCR-R4-MHC-I (noncognate) and LFA-1-ICAM-1 receptor-ligand bonds (**Fig. 6a)**. Consistent with the results in **Fig. 5**, as APC surface stiffness is increased, more elastic energy (in units of k_B_T) is stored in the TCR-ligand and adhesion bonds, with the greatest amount stored in LFA-1-ICAM-1 complexes, followed by OT1(†) TCR-OVA-MHC-I and OT1(†) TCR-R4-MHC-I complexes. But the elastic energy stored in the APC surface varies non-monotonically, reaching a peak near the optimal surface stiffness (∼1 pN/nm) followed by a rapid decrease at large stiffness values (> 100 pN/nm). This is consistent with the view of impedance matching; wherein soft APCs optimally transmit the deformation and force needed to improve TCR signal amplification.

**Fig. 6.**
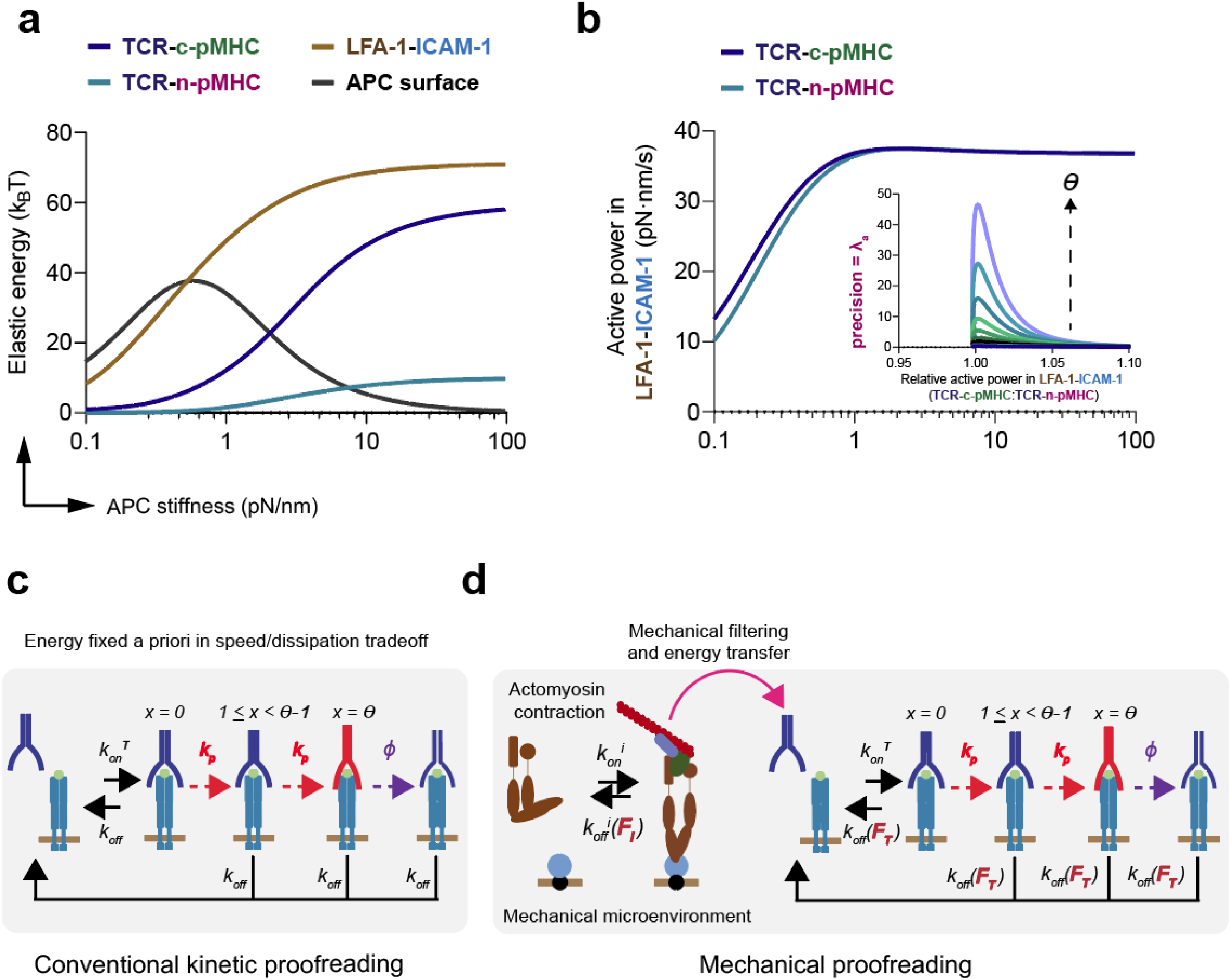
Mechanical filtering with LFA-1 determines TCR signal amplification in the mechanical proofreading framework. a) Elastic energy storage (in units of k_B_T) in TCR-pMHC bonds, LFA-1-ICAM-1 bonds, and the APC surface as a function of APC surface stiffness. b) Active power (energy transport) in total bound LFA-1-ICAM-1 complexes as a function of APC surface stiffness and TCR signal amplification precision as a function of the relative ratio of active power in LFA-1-ICAM-1 complexes when TCRs engage c-pMHC versus n-pMHC. c) Schematic of conventional kinetic proofreading mechanism in TCR-ligand discrimination, where the amount of free energy that is expended to improve TCR signaling precision is fixed a priori intracellularly in a speed/energy dissipation tradeoff. d) Schematic of mechanical proofreading mechanism in TCR-ligand discrimination involving cooperativity with LFA-1 coreceptors. In mechanical proofreading, Mechanical filtering between LFA-1-ICAM-1 bonds and the TCR determines mechanical energy transport from engaged contractile actomyosin filaments to the TCR to facilitate ligand proofreading precision.

In **Fig. 6b**, we also plot the active power dissipated by the collection of formed LFA-1-ICAM-1 complexes (,**eq. 25**) as a function of APC surface stiffness. We do not observe any significant differences in the active power dissipated by bound LFA-1-ICAM complexes when either OT1(†) TCR-OVA-MHC-I or OT1(†) TCR-R4-MHC-I bonds are engaged. Illustrating the proofreading precision per proofreading sequence as a function of the relative active power (TCR-c-pMHC:TCR-n-pMHC, **Fig. 6b inset**) further supports this, as the maximal precision is achieved around a relative power of unity (P_c_/P_n_ ∼ 1). Altogether, these results suggest that unlike conventional kinetic proofreading where a fixed amount of chemical energy is spent to increase precision and speed (involving a tradeoff with greater energy dissipation, **Fig. 6c**), our mechanokinetic framework decouples nonequilibrium dissipation from proofreading precision by employing a soft environment as a mechanical bottleneck that regulates the energization of discriminating reactions (**Fig. 6d**).

To further elucidate this, we devise a dimensionless scaling parameter 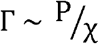 to compare the energy cost for nonequilibrium power generated by the actomyosin cortex and dissipated through the adhesive dynamics of the molecular clutch to the nonequilibrium chemical power dissipation dictated by the phosphorylation kinetics of the TCR. Here, the nonequilibrium chemical power dissipation is 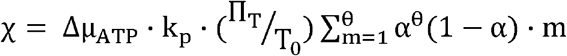 in which 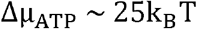 is the chemical potential associated with ATP hydrolysis and 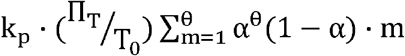 is the TCR phosphorylation flux. In **Appendix Fig. 4a**, we plot the average chemical energy necessary for TCR phosphorylation 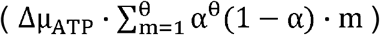 as a function of APC surface stiffness. Soft APC surfaces with intermediate stiffness ∼ 1 pN/nm allow for optimal TCR-c-pMHC phosphorylation whereas stiffnesses approaching 0 pN/nm or 1000 pN/nm leads to suboptimal chemical energy expenditures. This is additionally reflected in suppressed erroneous TCR-n-pMHC chemical energy expenditure for stiff substrates (σ_APC_ ≥ 1 pN/nm).

To evaluate the overall energetic efficiency of mechanical proofreading and impedance matching, we plot the dimensionless scaling parameter Γ as a function of normalized APC surface stiffness for OT1(†) TCR-OVA-MHC-I or OT1(†) TCR-R4-MHC-I (**Appendix Fig. 4b**). For noncognate OT1(†) TCR-R4-MHC-I slip bonds, the rapid collapse of χ in the denominator causes Γ to monotonically diverge as nonequilibrium power P increases with increasing APC surface stiffness. In this state, LFA-1-ICAM-1 catch bonds expend mechanical work but generates little TCR signaling output. Strikingly, for cognate OT1(†) TCR-OVA-MHC-I catch bonds, Γ exhibits a wave-like trajectory that maps the biophysical lifecycle of TCR-c-pMHC bonds under mechanical load. At low mechanical power P (norm. APC surface stiffness ≤ 1), mechanical work done by LFA-1 outpaces TCR signaling output, driving a local maximum in Γ. However, as mechanical power reaches the critical TCR-c-pMHC catch-bond threshold (norm. APC surface stiffness ∼ 1; 8-10 pN of applied force), a surge in chemical dissipation χ outpaces P, driving a local minimum in Γ, and thus realizing a ‘optimal’ mechanical proofreading regime in which the TCR extracts the maximal chemical signaling payout per unit of mechanical work invested by LFA-1 (e.g., ‘mechanical’ impedance matching). When mechanical power exceeds this optimum (norm. APC surface stiffness ≥ 1), the TCR-c-pMHC catch bond ruptures (entering the slip regime), χ minimizes, and Γ diverges upwards.

## Discussion

By integrating LFA-1 molecular clutch dynamics with a previously reported kinetic proofreading model with limited signaling, we developed a ‘mechanical’ proofreading framework of TCR-ligand discrimination, incorporating key features of endogenous TCR mechanosensation and its mechanical cooperativity with LFA-1 coreceptors. This minimal framework provides mechanistic support for how TCR/LFA-1 mechanical cooperation may directly energize the TCR-pMHC bond and enable force-sensitive ligand discrimination^10,16,18^. The integration of TCR-c-pMHC catch bonds or TCR-n-pMHC slip bonds into previously reported kinetic proofreading frameworks confirms that force may enhance proofreading precision by maximizing bond lifetime with c-pMHCs such that the probability of ‘correct’ TCR phosphorylation by relevant adaptor molecules and subsequent activation of signaling cascades downstream of the TCR occurs (**Fig. 1a-1e, Fig. 2b-2f**)^1^. Transient TCR-n-pMHC slip bonds are more likely to exit the phosphorylation pathway with respect to greater force application, thus minimizing proofreading error and erroneous activation.

These mechanical ‘checkpoints’ may explain how developing thymocytes are selected within the thymus through load-sensing through the Pre-TCR and TCR^59–61^, and how mature T-cells are able to be activated by as rare as a single c-pMHC molecule decorating an APC surface^62^. It is important to appreciate that though this may hold true for the αβTCR^9,63^, emerging evidence suggests that the γδTCR is force-agnostic^64,65^.

Energization of the TCR by actomyosin contractility through LFA-1-ICAM-1 adhesions is speculated to enhance TCR-ligand discrimination precision and stabilize the TCR-c-pMHC catch bond^18,66,67^. Our framework additionally supports this hypothesis through the prediction of physiological pN-level loading regimes on the TCR and LFA-1 in response to the mechanical properties of a model APC surface (**Appendix Fig. 2b, Fig. 4b-4e**)^68–70^. However, the role of actomyosin-rich T-cell microvilli protrusions^71,72^, the molecular crosstalk between TCR microcluster nucleation and LFA-1-ICAM-1 adhesions^44,45^, the complementary role of other co-effector molecules (e.g., CD4/CD8 and CD28)^73–80^, and the impact of extracellular matrix (ECM) mechanics^81–85^ on TCR-ligand discrimination remain significant points for further theoretical and experimental investigation.

Surprisingly, the mechanical cooperativity of LFA-1 coreceptors and the TCR in this framework does not follow a conventional kinetic proofreading mechanism that involves a precision/speed/energy dissipation tradeoff (**Fig. 6b, 6c**)^18,55–57^. Rather, Mechanical filtering between LFA-1 and the TCR mediated by the mechanical microenvironment determines the energy budget and how that energy is expended by the TCR to improve its performance in ligand discrimination (**Fig. 6d-f**)^63,86^. Thus, if the energy budget is indeed set by the microenvironment, it is likely that other topographical^87^ and physicochemical aspects of the mechanical microenvironment beyond T-cell-APC contact mechanics (e.g., the ECM^88,89^, lymphoid tissue microenvironment in both disease and homeostasis^90–92^, tumor microenvironments^9,93–96^, etc.) are important as well^97^.

The complex dynamics of TCR mechanical proofreading can be distilled into a simpler parametric framework governed by active power and mechanical coupling. In the asymptotic limits of substrate stiffness (approaching 0 pN/nm and 1000 pN/nm), the substrate effectively decouples the TCR and LFA-1 coreceptors. In these regimes, active energy transport is minimized, and the physical constraints on the TCR are dictated entirely by the passive properties of the membrane. However, at intermediate substrate stiffnesses, the substrate acts as a mechanical conduit. This allows the actomyosin-driven molecular clutch of LFA-1 to ‘talk’ to the TCR, actively tuning the force landscape to hit the catch/slip transition and maximize proofreading precision. To rigorously test and potentially falsify this model, future experimental strategies can actively decouple these parameters. For example, perturbing the intrinsic membrane stiffness via cholesterol doping^98^, or utilizing pMHC ligands with artificially high catch/slip transition forces^14,15,99,100^, should theoretically shift or abolish the proofreading optimum. By assessing TCR discriminatory precision against these modified parameters across both zero-stiffness and finite-stiffness substrates, experimentalists could directly validate the necessity of this mechanical circuit in active antigen discrimination.

Our findings emphasize the potential mechanical determinants of TCR signal amplification, and may have utility for exploring *in vitro* T-cell-material interface variable spaces and additional mechanoimmunological investigation of TCR-ligand discrimination, antigen recognition, and activation. The reported mathematical framework may additionally be used to dramatically constrain and formulate future mechanochemical feedback models of TCR signal transduction and T-cell activation.

## Methods

### Numerical methods

Nonlinear least-squares regression methods and simulations were implemented in MATLAB R2024a to solve constitutive equations or extract parameters from the literature (**Appendix Table a1**.). Data was visualized in either MATLAB R2024a or Prism v10.4.1.

## Acknowledgements

We thank Dr. Sungmin Nam (Harvard University), Dr. Yuesong Hu (Wyss Institute), Rohan Thakur (Massachusetts Institute of Technology), Dr. Kwasi Adu-Berchie (Wyss Institute), Dr. Andrew Khalil (Whitehead Institute and Wyss Institute), Dr. Vinny Chandran Suja (Harvard University), Dr. Debraj Ghose (Wyss Institute), Aditya Patil (Harvard University), Dr. Ross Jones (University of British Columbia), Dr. Ajinkya Ghagre (University of British Columbia), and Dr. Khalid Salaita (Emory University) for valuable scientific discussions and/or suggestions during the preparation of this manuscript. We thank Dr. Ze Gong (University of Science and Technology of China) for valuable scientific discussions on the initial theoretical concept and for providing example MATLAB implementations. We thank Dr. Melissa Lever (Marks and Clerk) for providing example MATLAB implementations. We acknowledge funding from the NIH NCI (Wyss Institute i3 center, F99/K00), Wellcome Leap Foundation (HOPE), and NSF (Harvard MRSEC).

## Appendix

Key model derivations, model parameters, parameter calculations and additional equations, and additional analytical data are provided in the appendix. MATLAB script of the model implementation encompassing all master equations are uploaded on <u>GitHub</u>.

## Author contributions

N.J. and S.S. (concept, theory, implementation, data visualization, analysis, writing). J.M.B and T.H. (concept, theory, implementation). B.A.N. and W.H.J. (implementation). P.W.Z. (writing). L.M. and D.J.M. (concept, theory, writing).

