## Appendix for "Substrate mediated mechanical forces enable optimal kinetic proofreading by T-cell receptors"

**Additional analytical data**

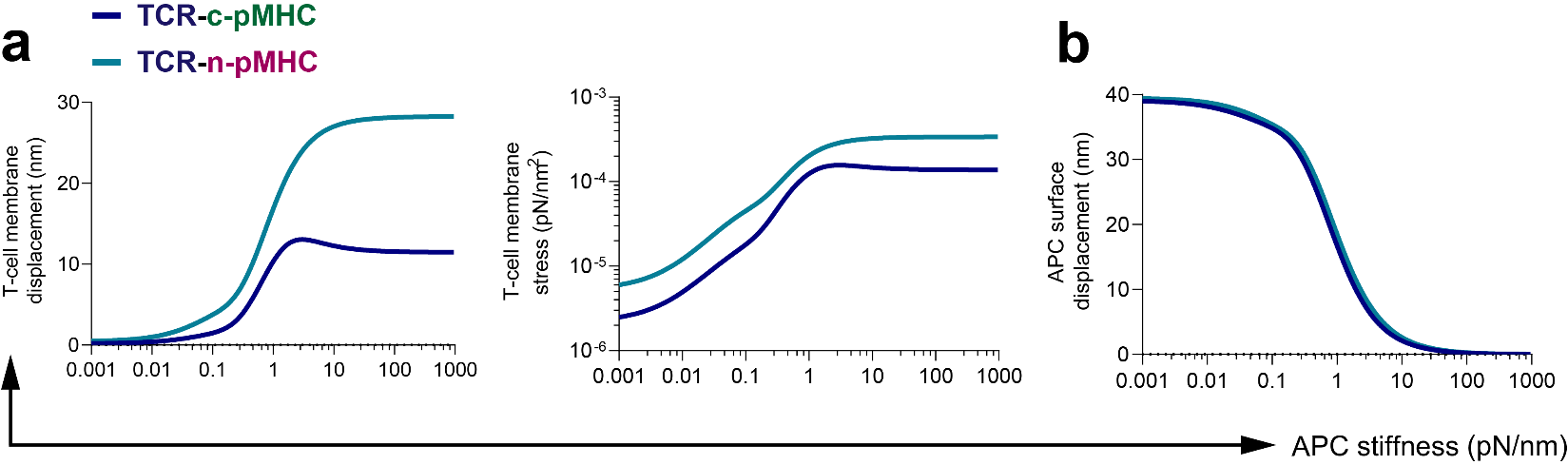
 **Appendix Fig. 1. T-cell membrane mechanics and APC surface displacement as a function of APC stiffness.** a) T-cell membrane displacements and membrane stresses when TCRs are engaged with either c-pMHCs or n-pMHCs. b) APC surface displacements for TCR-c-pMHCs or TCR-n-pMHCs.

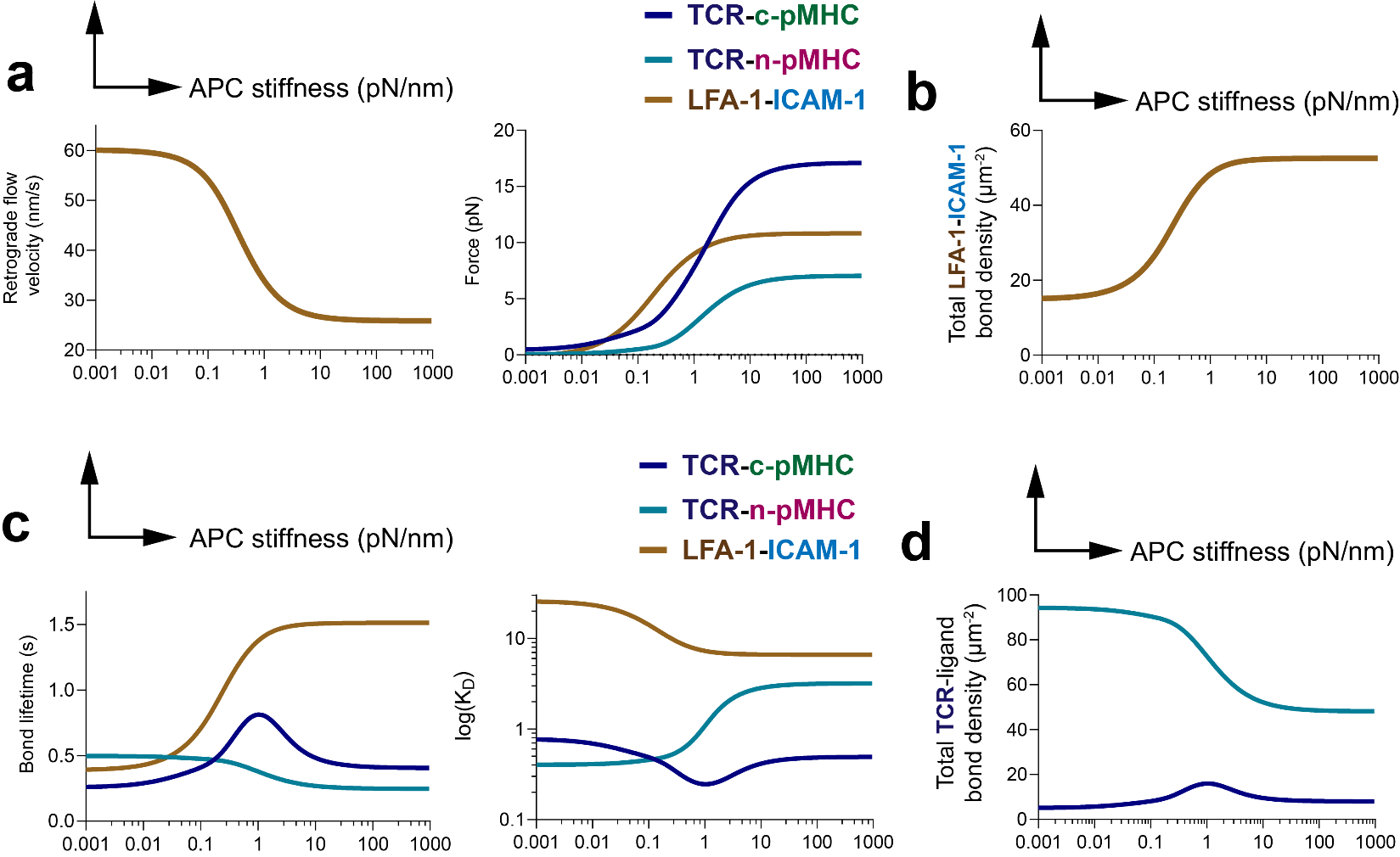

**Appendix Fig. 2. LFA-1 molecular clutch dynamics at steady-state modulates receptor-ligand force magnitude and bond formation kinetics in response to APC surface stiffness.** a) Actomyosin retrograde flow velocity and receptor generated forces as a function of APC stiffness. b) Total LFA-1-ICAM-1 bond density as a function of APC stiffness. c) Bond lifetimes and dissociation constants of TCR-c-pMHC, TCR-n-pMHC, and LFA-1-ICAM-1 as a function of APC stiffness. d) TCR-ligand bond interfacial surface densities as a function of APC stiffness.

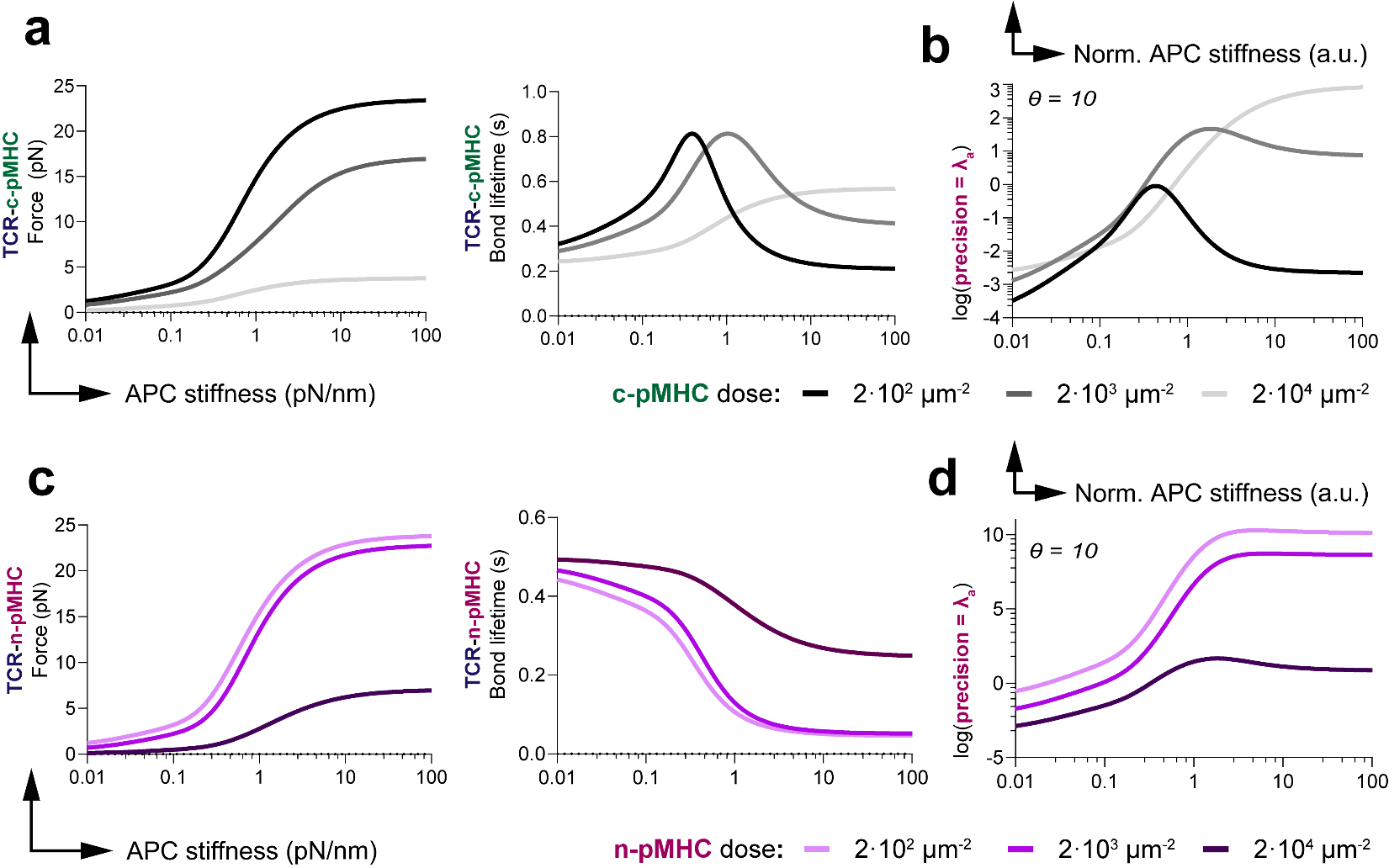

**Appendix Fig. 3. TCR-generated forces, TCR-pMHC bond lifetimes, and proofreading precision as a function of APC stiffness and pMHC dose.** a) TCR-c-pMHC generated forces and bond lifetimes with differing c-pMHC dose. b) Proofreading precision with differing c-pMHC dose after θ = 10 proofreading sequences. c) TCR-n-pMHC generated forces and bond lifetimes with differing n-pMHC dose. d) Proofreading precision with differing n-pMHC doses after θ = 10 proofreading sequences. When c-pMHC dose is changed, n-pMHC is fixed to 2·10^4^ µm^-2^ whereas when n-pMHC dose is changed, c-pMHC is fixed to 2·10^3^ µm^-2^. APC stiffness is normalized to 1 pN/nm.

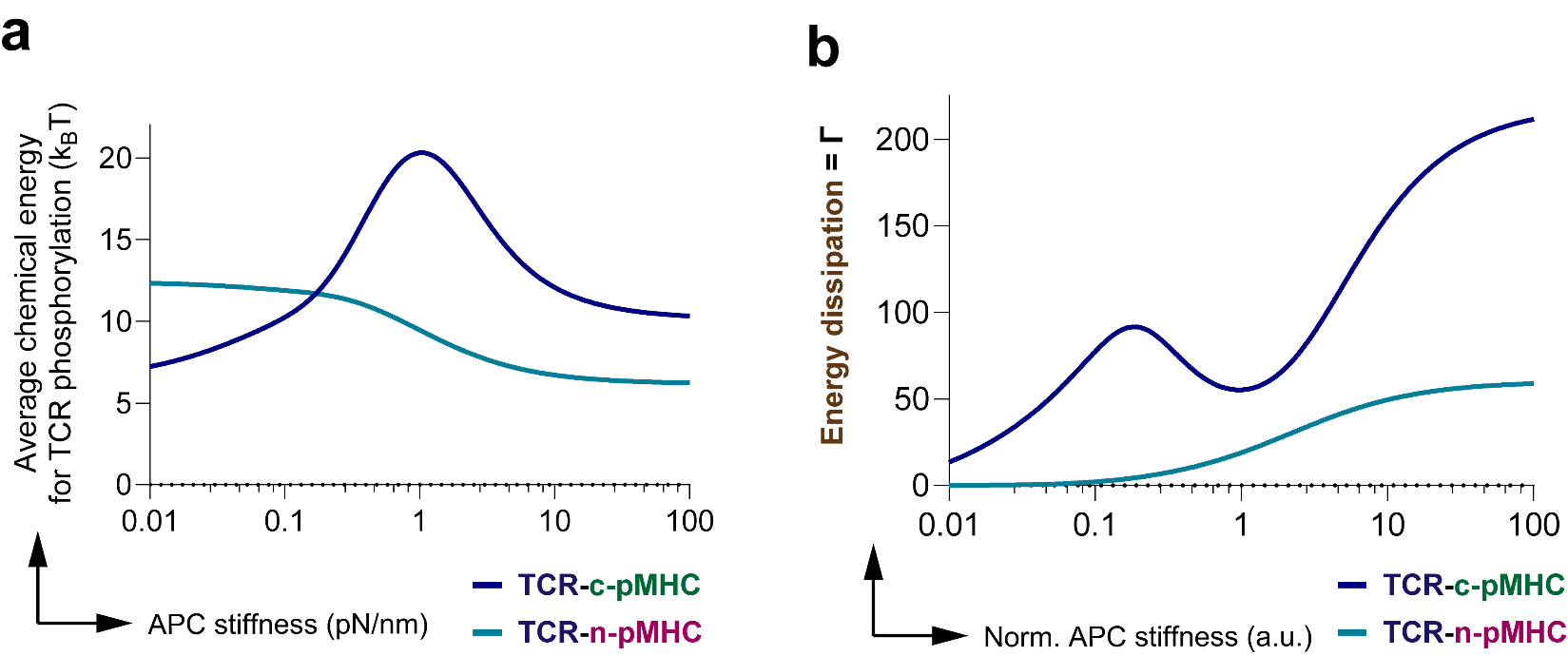

**Appendix Fig. 4. Average chemical energy for TCR phosphorylation and energetic efficiency as a function of APC stiffness.** a) Average chemical energy for TCR phosphorylation in TCR-pMHC bonds as a function of APC surface stiffness. b) Dimensionless scaling parameter illustrating the energetic efficiency of the mechanical proofreading framework as a function of dimensionless APC surface stiffness. APC surface stiffness is normalized to 1 pN/nm.

**Model derivations**

Derivations of **eq. 4** and **eq. 5**

The steady-state solutions for the mass action kinetic rate laws describing their respective, force-sensitive biochemical processes for formation of bound TCR-pMHC complexes is admitted by invoking the principle of mass conservation, using a previously reported model for kinetic proofreading with limited signaling^1^. The governing system of ordinary differential equations (ODEs) for this scheme are (**eq. a1-a6**):

**(a1)**

$$\frac{\mathrm{dp}}{\mathrm{dt}}={-k}_{\mathrm{on}}^{T}\cdot pT+k_{\mathrm{off}}\left( F_{T} \right)\cdot\left[ \sum_{x=0}^{\theta} \Pi_{x}+\Pi_{\theta}^{-} \right]$$

**(a2)**

$$\frac{dT}{\mathrm{dt}}={-k}_{\mathrm{on}}^{T}\cdot pT+k_{\mathrm{off}}(F_{T})\cdot\left[ \sum_{x=0}^{\theta} \Pi_{x}+\Pi_{\theta}^{-} \right]$$

**(a3)**

$$\frac{d\Pi_{0}}{\mathrm{dt}}=k_{\mathrm{on}}^{T}\cdot pT {- (k_{p}+k}_{\mathrm{off}}(F_{T}))\cdot\Pi_{0}$$

**(a4)**

$\frac{d\Pi_{x}}{\mathrm{dt}}=k_{p}\cdot\Pi_{x-1}{- (k_{p}+k}_{\mathrm{off}}(F_{T}))\cdot\Pi_{x}$ for $1\leq x< \theta-1$

**(a5)**

$$\frac{d\Pi_{\theta}}{\mathrm{dt}}=k_{p}\cdot\Pi_{\theta-1}{- (\phi+k}_{\mathrm{off}}(F_{T}))\cdot\Pi_{\theta}$$

**(a6)**

$$\frac{d\Pi_{\theta}^{-}}{\mathrm{dt}}=\phi\cdot\Pi_{\theta}{- k}_{\mathrm{off}}(F_{T})\cdot\Pi_{\theta}^{-}$$

where $p$ is the unbound pMHC dose (c or n-pMHC), $T$ is the unbound T-cell membrane density of TCRs in the contact area, $\Pi_{\theta}$ is the interfacial surface density of activated signaling-competent bound TCR-pMHC complexes after $\theta$ proofreading steps at rate $k_{p}$, $\Pi_{\theta}^{-}$ is the interfacial surface density of TCR-pMHC complexes that have been rendered non-signaling at a decay rate $\phi$, and$k_{\mathrm{off}}(F_{T})$ is the force-sensitive TCR-pMHC reverse reaction rate. The total interfacial surface density of bound TCR-pMHC complexes $\Pi_{T}$ (either $\Pi_{c}$ or $\Pi_{n}$) is taken from **(a1)** and **(a2)** (**eq. a8**):

**(a8)**

$$\Pi_{T}=\sum_{x=0}^{\theta} \Pi_{x}+ \Pi_{\theta}^{-}$$

The net unbound pMHC dose $p_{0}$ (either $p_{c}$ or $p_{n}$) and net unbound TCR membrane density $T_{0}$ are conserved quantities defined by the following mass conservation equations (**eq. a9** and **eq. a10**):

**(a9)**

$$p_{0}=p+\Pi_{T}$$

**(a10)**

$$T_{0}=T+\Pi_{T}$$

substituting **(a9)**, **(a10)**, and the force-sensitive dissociation constant $K_{D}= {k_{\mathrm{off}}(F_{T})}/{k_{\mathrm{on}}^{T}}$ into the governing system of ODEs at steady-state admits the solution for the interfacial surface density for total bound TCR-pMHC complexes (**eq. 4** and **eq. 5**):

$$\Pi_{T}=\frac{1}{2}\left( p_{0}+T_{0}+K_{D} \right)-\frac{1}{2}\sqrt{\left( p_{0}+T_{0}+K_{D} \right)^{2}-4p_{0}T_{0}}$$

Derivations of **eq. 6-9**

The total interfacial surface density of bound TCR-pMHC complexes at steady-state **(a8)** can be rewritten as (**eq. a11**):

**(a11)**

$$\Pi_{T}=\sum_{x=0}^{\theta-1} \Pi_{x}+\Pi_{\theta}+ \Pi_{\theta}^{-}$$

solving for$\Pi_{x}$ and $\Pi_{\theta}^{-}$ from the system of governing ODEs at steady-state and then substituting into **(a11)** yields (**eq. a12**):

**(a12)**

$$\Pi_{T}=\frac{1-\left( \frac{k_{P}}{k_{p}+k_{\mathrm{off}}(F_{T})} \right)^{\theta}}{1-\frac{k_{p}}{k_{p}+k_{\mathrm{off}}(F_{T})}}\cdot\Pi_{0} {+ \Pi}_{\theta}+\frac{\phi}{k_{\mathrm{off}}(F_{T})}\cdot\Pi_{\theta}$$

**eq. 8** and **eq. 9** can then be defined from **(a12)** as:

$$\alpha=\frac{k_{P}}{k_{p}+k_{\mathrm{off}}(F_{T})}$$

By equating **(a2)** with **(a3)** at steady-state, $\Pi_{T}$ can be substituted with (**eq. a13**):

**(a13)**

$$\Pi_{T}=\frac{\Pi_{0}}{1-\alpha}$$

substitution of **(a13)** into **(a12)** yields the function for **eq. 6** and **eq. 7**:

$$\Pi_{\theta}=\frac{k_{\mathrm{off}}(F_{T})}{k_{\mathrm{off}}(F_{T})+\phi}\cdot\alpha^{\theta}\Pi_{T}$$

Derivation of **eq. 3 and eq. 14-19**

To describe the molecular clutch behavior of LFA-1 adhesion molecules, a stochastic model for LFA-1-ICAM-1 adhesion kinetics is described. A bound LFA-1-ICAM-1 complex is asserted to behave as a Hookean spring such that the length of the full complex is $l_{i}=h+y$ where $h$ is the interfacial distance between the T cell and APC membranes, and the elastic force on an LFA-1-ICAM-1 bond is $F_{i}= \kappa_{i}\cdot\left( l_{i}-l_{i}^{0} \right)$. The probability of a bound LFA-1-ICAM-1 complex at time $t$ with a stretch length $y$ is $P(y, t)$ which obeys the following chemical master equation (**eq. a14**):

**(a14)**

$$\frac{\partial P}{\partial t}+\frac{\partial}{\partial y}\left( P\cdot\frac{\partial y}{\partial t} \right)=k_{\mathrm{on}}^{i}\cdot\left( 1-n(t) \right)\delta\left( y \right)-k_{\mathrm{off}}^{i}\left( F_{i} \right)\cdot P$$

where $n\left( t \right)= \int P\left( y, t \right)\mathrm{dy}$ such that $1-n\left( t \right)$ is the probability of LFA-1 being unbound and $\delta\left( y \right)$ is the Kronecker-delta function. LFA-1 is assumed to bind with zero stretch ($y=0$) but after binding to ICAM-1, the complex stretches due to retrograde flow of the actomyosin gel given by ${\partial y}/{\partial t}=v_{a}$. Rather than working within the full probability distribution, it is beneficial to work within its low order moments; the bound fraction $n\left( t \right)$ and average LFA-1-ICAM-1 stretch length $\left\langle y(t) \right\rangle=\frac{1}{n(t)}\int y\cdot P\left( y, t \right)\mathrm{dy}$. Neglecting all higher order fluctuations/moments (akin to a mean-field approximation), the first two moments of **(a14)** are taken to obtain (**eq. a15**) and (**eq. a16**):

**(a15)**

$$\frac{\partial n}{\partial t}=k_{\mathrm{on}}^{i}\cdot\left( 1-n\left( t \right) \right)-k_{\mathrm{off}}^{i}(F_{i})\cdot n\left( t \right)$$

**(a16)**

$$\frac{\partial\left\langle y_{i} \right\rangle}{\partial t}=v_{a}-k_{\mathrm{on}}^{i}\cdot\left( \frac{1}{n(t)}-1 \right) \left\langle y_{i} \right\rangle$$

Consistent with the mean-field approximation, the averages of all nonlinear terms have been factored by neglecting correlations (e.g., $k_{\mathrm{on}}^{i}\approx k_{\mathrm{off}}^{i}(F_{i})$). Imposing $i\left( t \right)=i_{0}\cdot n\left( t \right)$, **eq. 14** and **eq. 16** are obtained:

$$\frac{di(t)}{\mathrm{dt}}=k_{\mathrm{on}}^{i}\cdot\left( i_{0}-i(t) \right)-k_{\mathrm{off}}^{i}(F_{i})\cdot i(t)$$

$$\frac{d\left\langle y_{i} \right\rangle}{\mathrm{dt}}=v_{a}-k_{\mathrm{on}}^{i}\cdot\left( \frac{i_{0}}{i\left( t \right)}-1 \right) \left\langle y_{i} \right\rangle$$

Model parameters, parameter calculations, and relevant equations

**Table a1.** Main model parameters

| **Parameter** | **Symbol** | **Value** | **Unit** | **Reference: DOI (location) [or calculated]** |
| --- | --- | --- | --- | --- |
| TCR-pMHC bond stiffness | $\kappa_{T}$ | 0.6 | pN/nm | doi.org/10.1038/s41467-022-34587-w  (Fig. 5C)^2^ |
| LFA-1-ICAM-1 bond stiffness | $\kappa_{i}$ | 0.2 | pN/nm | doi.org/10.1371/journal.pcbi.1011237^3^ |
| LFA-1-ICAM-1 on rate | $k_{\mathrm{on}}^{i}$ | 0.1 | sec^-1^ | doi.org/10.1371/journal.pcbi.1004481 (Table 1)^4^ |
| LFA-1-ICAM-1 catch rate | $k_{c,i}^{0}$ | 2.5 | sec^-1^ | doi.org/10.1074/jbc.m110.155770 (Least-squares fit from Fig. 1A of bond-lifetime for Ca^2+^/Mg^2+^/CXCL12 condition)^5^ |
| LFA-1-ICAM slip rate | $k_{s,i}^{0}$ | 1.4·10^-2^ | sec^-1^ | doi.org/10.1074/jbc.m110.155770 (Least-squares fit from Fig. 1A of bond-lifetime for Ca^2+^/Mg^2+^/CXCL12 condition)^5^ |
| LFA-1-ICAM-1 catch force | $F_{c,i}^{0}$ | 6.2 | pN | doi.org/10.1074/jbc.m110.155770 (Least-squares fit from Fig. 1A of bond-lifetime for Ca^2+^/Mg^2+^/CXCL12 condition)^5^ |
| LFA-1-ICAM-1 slip force | $F_{s,i}^{0}$ | 3.9 | pN | doi.org/10.1074/jbc.m110.155770 (Least-squares fit from Fig. 1A of bond-lifetime for Ca^2+^/Mg^2+^/CXCL12 condition)^5^ |
| TCR on rate (per bond) | $k_{\mathrm{on}}^{T}$ | 5.0·10^-6^ | μm^2^sec^-1^ | doi.org/10.1038/nri3728 (supplementary info), Calculated by dividing by TCR membrane density^1^ |
| TCR-c-pMHC catch rate | $k_{c,c}^{0}$ | 3.99 | sec^-1^ | doi.org/10.1038/s41467-023-38267-1 (supplementary info, Table 4, OT1(†) /OVA/H2-Kb (α3A2/hβ2m)) - TCR/peptide/MHC-I^6^ |
| TCR-c-pMHC slip rate | $k_{s,c}^{0}$ | 0.41 | sec^-1^ | doi.org/10.1038/s41467-023-38267-1 (supplementary info, Table 4, OT1(†) /OVA/H2-Kb (α3A2/hβ2m)) – TCR/peptide/MHC-I^6^ |
| TCR-c-pMHC catch force | $F_{c,c}^{0}$ | 3.03 | pN | doi.org/10.1038/s41467-023-38267-1 (supplementary info, Table 4, OT1(†) /OVA/H2-Kb (α3A2/hβ2m)) - TCR/peptide/MHC-I, Calculated (**eq. a17**)^6^ |
| TCR-c-pMHC slip force | $F_{s,c}^{0}$ | 9.57 | pN | doi.org/10.1038/s41467-023-38267-1 (supplementary info, Table 4, OT1(†) /OVA/H2-Kb (α3A2/hβ2m)) – TCR/peptide/MHC-I, Calculated (**eq. a17**)^6^ |
| TCR-n-pMHC slip rate | $k_{s,n}^{0}$ | 2.0 | sec^-1^ | doi.org/10.1038/s41467-023-38267-1 (supplementary info, Table 4, OT1(†) /R4/H2-Kb (α3A2/hβ2m)) – TCR/peptide/MHC-I^6^ |
| TCR-n-pMHC slip force | $F_{s,n}^{0}$ | 10 | pN | doi.org/10.1038/s41467-023-38267-1 (supplementary info, Table 4, OT1(†) /R4/H2-Kb (α3A2/hβ2m)) – TCR/peptide/MHC-I, Calculated (**eq. a17**)^6^ |
| TCR membrane density | $T_{0}$ | 2000 | μm^-2^ | doi.org/10.1073/pnas.1605399113^7^ |
| LFA-1 membrane density | $i_{0}$ | 400 | μm^-2^ | doi.org/10.1371/journal.pcbi.1004481 (Table 1)^4^ |
| c-pMHC surface density | $p_{c}$ | 2000 | μm^-2^ | N.A. |
| n-pMHC surface density | $p_{n}$ | 2·10^4^ | μm^-2^ | N.A. |
| Surface stiffness | $\sigma_{\mathrm{APC}}$ | Free parameter | pN/nm | N.A. |
| T-cell membrane surface tension | $\gamma$ | 3·10^-2^ | pN/nm | doi.org/10.1016/j.sbi.2015.07.010^8^ |
| Active stress in actomyosin gel | $\sigma_{a}$ | 0.001 | pN/nm^2^ | doi.org/10.1038/nphys3224, Calculated (**eq. a18**)^9^ |
| Actomyosin gel unloaded retrograde flow velocity | $v_{0}$ | 60 | nm/sec | doi.org/10.1038/ncb3191^10^ |
| Equilibrium separation | $h_{0}$ | 55 | nm | doi.org/10.1038/s42003-021-02995-1 (approximate length of CD45 ectodomain)^11^ |
| TCR-pMHC complex bond length | $l_{T}^{0}$ | 15 | nm | doi.org/10.1371/journal.pcbi.1004481 (Table 1)^4^ |
| LFA-1-ICAM-1 complex bond length | $l_{i}^{0}$ | 40 | nm | doi.org/10.1371/journal.pcbi.1004481 (Table 1)^4^ |
| Number of proofreading sequences | $\theta$ | 10 | A.U. | doi.org/10.1038/nri3728 (supplementary info)^1^ |
| Tyrosine phosphorylation rate | $k_{p}$ | 1 | sec^-1^ | doi.org/10.1038/nri3728 (supplementary info)^1^ |
| TCR-pMHC signaling complex decay rate | $\phi$ | 0.1 | sec^-1^ | doi.org/10.1038/nri3728 (supplementary info)^1^ |
| TCR microcluster radii | $R$ | 50 | nm | doi.org/10.1111%2Fimr.12120^12^ |
| Contact area | $A_{c}$ | 3.0·10^4^ | nm^2^ | doi.org/10.1073/pnas.1605399113^7^ |

**Table a2.** Additional model parameters for OT1(†) TCR-c-pMHC binding

| **Parameter** | $k_{c,c}^{0}$ (sec^-1^) | $k_{s,c}^{0}$ (sec^-1^) | $F_{c,c}^{0}$ (pN) | $F_{s,c}^{0}$ (pN) | **Reference: DOI (location) [or calculated]** |
| --- | --- | --- | --- | --- | --- |
| A2 antigen | 3.36 | 0.86 | 4.07 | 11.8 | doi.org/10.1038/s41467-023-38267-1 (supplementary info, Table 4, OT1(†) /A2/H2-Kb (α3A2/hβ2m)) – TCR/peptide/MHC-I, Calculated (**eq. a17**)^6^ |
| G4 antigen | 1.99 | 1.47 | 1.31 | 3.96 | doi.org/10.1038/s41467-023-38267-1 (supplementary info, Table 4, OT1(†) /G4/H2-Kb (α3A2/hβ2m)) – TCR/peptide/MHC-I, Calculated (**eq. a17**)^6^ |
| E1 antigen | 0.39 | 2.01 | 1.77 | 4.33 | doi.org/10.1038/s41467-023-38267-1 (supplementary info, Table 4, OT1(†) /E1/H2-Kb (α3A2/hβ2m)) – TCR/peptide/MHC-I, Calculated (**eq. a17**)^6^ |

Set of additional equations for parameter and thermodynamic calculations:

The characteristic catch/slip transition state forces for TCR binding of either c-pMHC or n-pMHC were calculated using the following equation after characteristic transition state displacements were extracted (**eq. a17**):

**(a17)**

$$F_{\neq}^{0}=\frac{k_{B}T}{\delta_{\neq}^{0}}$$

where $F_{\neq}^{0}$ is the characteristic transition state force for either catch or slip transitioning, $\delta_{\neq}^{0}$ is the transition state distance, and $k_{b}T$ is thermal energy (~4.114 pN·nm). The active stress $\sigma_{a}$ in the actomyosin gel is calculated using the following equation (**eq. a18**):

**(a18)**

$$\sigma_{a}=\frac{n_{m}\cdot F_{m}}{l_{m}^{2}}$$

in which $n_{m}$ is the number of myosin motors, $F_{m}$ is the myosin-generated stalling force, and $l_{m}$ is the approximate size of a myosin mini-filament.

The standard state change in Gibbs free energy $\Delta G^{^{\circ}}$ for force-sensitive TCR-pMHC binding is calculated via the following relationship (**eq. a19**):

**(a19)**

$$\Delta G^{^{\circ}}=k_{B}T\cdot\ln(K_{D})$$

where $k_{B}T$ is thermal energy and $K_{D}={k_{\mathrm{off}}(F_{T})}/{k_{\mathrm{on}}^{T}}$ is the force-sensitive dissociation constant.
